# Quantitative assessment of cell population diversity in single-cell landscapes

**DOI:** 10.1101/333393

**Authors:** Qi Liu, Charles A. Herring, Quanhu Sheng, Jie Ping, Alan J. Simmons, Bob Chen, Amrita Banerjee, Guoqiang Gu, Robert J. Coffey, Yu Shyr, Ken S. Lau

**Affiliations:** Department of Biostatistics, Vanderbilt University Medical Center, Nashville, TN 37232; Center for Quantitative Sciences, Vanderbilt University Medical Center, Nashville, TN 37232; Epithelial Biology Center, Vanderbilt University Medical Center, Nashville, TN 37232; Program in Chemical and Physical Biology, Vanderbilt University School of Medicine, Nashville, TN 37232; Department of Cell and Developmental Biology, Vanderbilt University School of Medicine, Nashville, TN 37232; Department of Medicine, Vanderbilt University Medical Center, Nashville, TN 37232; Veterans Affairs Medical Center, Tennessee Valley Healthcare System, Nashville, TN 37232

## Abstract

Single-cell RNA-sequencing (scRNA-seq) has become a powerful tool for the systematic investigation of cellular diversity. As a number of computational tools have been developed to identify and visualize cell populations within a single scRNA-seq dataset, there is a need for methods to quantitatively and statistically define proportional shifts in cell population structures across datasets, such expansion or shrinkage, or emergence or disappearance of cell populations. Here we present sc-UniFrac, a framework to statistically quantify compositional diversity in cell populations between single-cell transcriptome landscapes. sc-UniFrac enables sensitive and robust quantification in simulated and experimental datasets in terms of both population identity and quantity. We have demonstrated the utility of sc-UniFrac in multiple applications, including assessment of biological and technical replicates, classification of tissue phenotypes, identification and definition of altered cell populations, and benchmarking batch correction tools. sc-UniFrac provides a framework for quantifying diversity or alterations in cell populations across conditions, and has broad utility for gaining insight on how cell populations respond to perturbations.

## Introduction

Single-cell sequencing technologies enables profiling of hundreds to thousands individual cells from a tissue comprising of a diversity of cell types [1–8]. The rapid advances of single-cell technologies have led to a proliferation of novel computational tools. Current analyses mainly focus on the identification of cell populations and transitional trajectories from data obtained from a single sample (reviewed in [9–12]). While these tools can shed light on complex biological processes within a given sample [13], it is emerging that the power of single-cell technologies lies in multi-sample experimental designs where responses of single-cells, in terms of identity and quantity, under multiple conditions can be assessed. This poses a new statistical challenge: given that each sample is comprised of transcriptomes from hundreds to thousands of individual cells, how does one compare across different samples to statistically assess population diversity and to detect population changes in an unbiased way? As increasing numbers of studies are generating scRNA-seq over multiple samples, there is an unmet need for a statistical framework that enables quantitative comparisons across single-cell landscapes. The framework would have wide application, from quantitatively uncovering cell population changes, assessing batch effect correction methods, to classifying disease subtypes based upon single-cell landscapes.

There currently exists a paucity of approaches aiming to determine cellular composition similarities and differences between samples. CITRUS is a supervised approach for identifying cell populations that are significantly different between specified outcomes, whose goal is distinct from unsupervised comparison of samples based on similarities [14]. Another approach uses the Wasserstein metric known as Earth mover’s Distance (EMD), which is a measure of the distance between two probability distributions over a certain data space [15]. Briefly, EMD partitions the entire data space into bins and measures the cost of transfer of data points from one distribution over these bins to resemble the other distribution. Orlova et al. applied EMD across datasets to quantify the similarity of two cell populations by measuring the distance between expression distributions in 2-dimensional marker space [16]. However, partition of data spaces into bins has an exponential computational cost as the number of dimensions increases, limiting EMD use to 1- or 2-dimensional data spaces. Although there are multiple dimension-reduction approaches for high dimensional data [11,17,18], analysis in a customizable, unrestricted number of dimensions would be preferred, especially for scRNA-seq datasets with thousands of native dimensions. We have developed the p-Creode score, which determines the similarities between p-Creode trajectories derived from a multi-dimensional single-cell landscape [19]. However, this approach is limited to datasets where trajectories can be derived from continuous single-cell data and is not generalizable to all data distributions. Currently, the most common strategy for assessing similarities between single-cell landscapes remains a visual evaluation of the degree of “mixing” of data points when two or more samples are analyzed together on a t-SNE plot [2]. Two methods have recently been developed to use k-nearest neighbors to characterize this degree of intermixing [20,21]. Both methods use the simple assumption that k-nearest neighbors of each cell should have the same distribution of sample labels as the full dataset if the datasets are well-mixed.

We were originally inspired by some of the early single-cell work where similarities between replicates can be qualitatively evaluated by the degree of mixing of hierarchical clusters formed between replicates [7]. Thus, by deriving a quantitative measure to compare between hierarchical trees generated by clustering, we can obtain a corresponding quantitative, statistically testable metric to compare cell population diversity between single-cell landscapes. Multiple metrics to measure similarity between trees have been proposed such as, Baker’s gamma index [22], which we have used previously to determine the similarity between signaling modules [23]. UniFrac is a distance metric originally devised to utilize phylogenetic tree comparisons to determine similarity in microbial communities between two ecologies [24]. UniFrac determines the similarity between trees by statistically evaluating shared branches that propagate through the entire structure of the data. UniFrac enables a statistically robust framework, where the weight of branches can be directly related to the number of organisms belonging to phylogenetic groups. We modified UniFrac to what we called sc-UniFrac (single-cell UniFrac) such that single-cell landscape comparisons can directly leverage the entire statistical framework built around UniFrac. sc-UniFrac compares diversity based on transcriptome similarities of single cells, and is more powerful than intermixing methods due to its account of entire data structures of the landscape. We demonstrated the utility of sc-UniFrac on quantifying similarities between simulated and real sc-RNAseq datasets, where the ground truth of similarities between samples is known. We also successfully applied sc-UniFrac on assessing the performance of single-cell batch-correcting methods. We envision that quantitative metrics such as sc-UniFrac will find increasing utility as larger volumes of sc-RNAseq data over many samples are generated. sc-UniFrac will greatly facilitate single-cell studies, include those aimed at deciphering how cell populations respond to perturbations or tracking the evolution of cell populations during disease progression.

## Results

### Overview of sc-UniFrac

For quantifying cell diversity differences in single-cell landscapes, we borrowed the UniFrac concept from the field of microbiome research. UniFrac is a distance metric for quantifying differences in phylogenetic diversity between ecological landscapes. Instead of operating on trees constructed on phylogenetic dissimilarities between two samples, sc-UniFrac analogously builds hierarchical trees from clustering transcriptional profiles of singles cells combined from two samples. Note that the purpose of the clustering is not to define cell populations within the samples but to partition the data points in a reproducible way. sc-UniFrac reconstructs a hierarchical tree from k cluster centroids independent of the clustering method used. The clustering tree, encompassing the structural features of cell subpopulations, is used for calculating the weighted UniFrac distance by assign values to each branch based on the relative abundance of cells (**Figure 1A**). Statistical significance of the distance between two samples is determined within the standard UniFrac framework, by permuting the sample labels of each cell on the tree without changing the tree structure. In this way, a distribution of distance is obtained with a p-value that reflects the probability that the permuted distances are greater than or equal to the observed distance by chance **(Methods)**. sc-UniFrac further identifies significantly altered cell populations that drive the compositional difference between single-cell transcriptome landscapes and detects gene signatures to mark these cell populations. Finally, sc-UniFrac predicts the potential identities of these populations by matching individual cell signatures to cell types from reference atlases. The general workflow of the sc-UniFrac pipeline is shown in **Figure 1B**.

**Figure 1:**
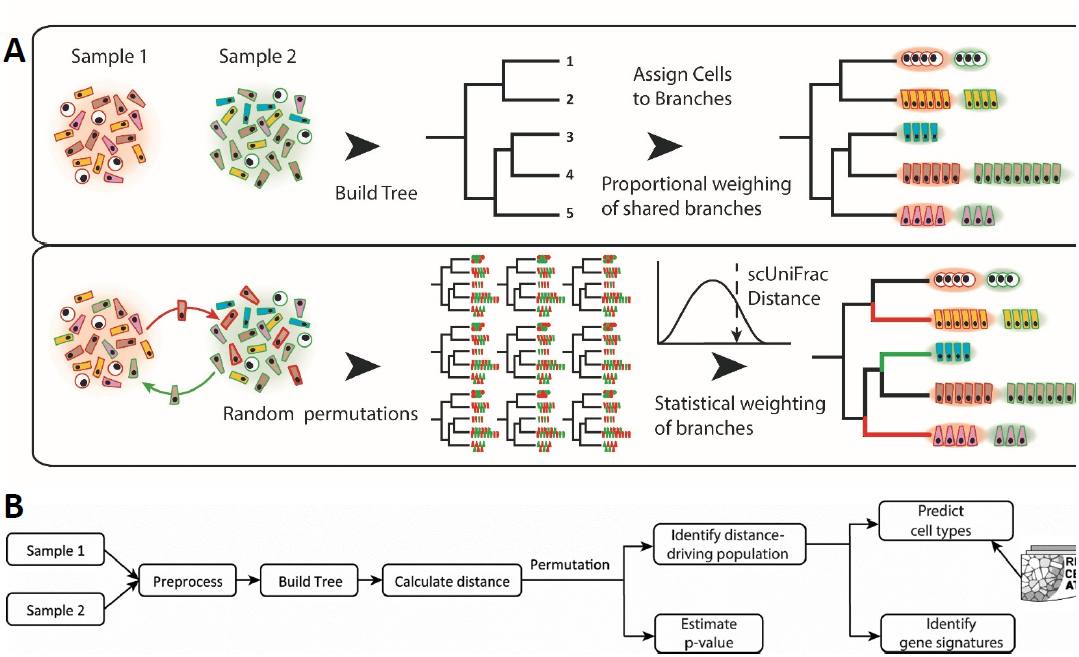
Overview of the sc-UniFrac method. (A) Distances in data space between single cells from 2 samples were used to build a common hierarchical tree. Each cell was then assigned to branches where proportional weighing of shared branches is used to calculate a UniFrac distance. In the second step, the labels between samples were randomized to generate a distribution of permuted distances, where a p-value for the sc-UniFrac distance can be calculated. (B) Workflow overview of the sc-UniFrac package for characterizing dissimilarities between samples.

### Sensitive and robust quantification of proportional shifts population diversity in single-cell landscapes

To evaluate the performance of sc-UniFrac on quantifying the compositional difference between single-cell landscapes, we first applied the method on experimental datasets where population structure can be precisely controlled. Two 1000-cell populations were generated by sampling from the CD8 and CD4 T cell populations, respectively, from the T-cell development dataset of the mouse thymus [25]. This process was repeated 50 times to evaluate the robustness of sc-UniFrac. Comparing the CD8 versus CD4 populations (1000-cell sample from each population) using sc-UniFrac revealed, as expected, they are completely different in their population structures with a large distance **(Figure 2A – red arrow, S1A)**. In contrast, comparing two identical 1000-cell populations resampled from CD8 cells revealed that they possess the same population structures with a median distance of 0 **(Figure 2A – green arrow, S1B)**. We then evaluated the performance of sc-UniFrac in a simulation experiment where we constructed a series of paired samples with a gradation of proportional shift in cell populations. For each pair, one sample included only CD8 cells (N1), while the other was comprised of proportional mixtures of CD4 and CD8 cells (N2), starting from 0% of CD4 cells (no shift) to 100% of CD4 cells (complete shift). The distance of sc-UniFrac progressively increased as proportional shifts became larger **(Figure 2A, S1C-D)**. Among the 50 resampled runs, less than 0.2% of the sc-UniFrac distances generated were significantly different (identified as false positives) when two samples were identical (0% proportional shift), while over 95% of the distances were significantly different even when there was as little as 2% of CD4 cells mixed in with the CD8 cells **(Figure 2B)**. These results signify that sc-UniFrac is sensitive and specific for detecting minute shifts in population structure.

Next, using this controlled sampling scheme, we evaluated the impact of changing various parameters on the performance of sc-UniFrac. First, we altered the k parameter for dividing the data into k sub-populations for analysis, which tunes the resolution by which sc-UniFrac analyzes the datasets. sc-UniFrac, regarding its quantitative ability and its sensitivity, was robust as long as k was not exceedingly low (k>3) **(Figure 2C, D)**. At k<=3, single-cell datasets are represented only by three major groups or less, reducing the resolution of sc-UniFrac for detecting differences in single-cell level diversity. Second, we assessed the effect of imbalanced dataset sizes. Keeping the size of sample 1 (N1) constant at 1000 cells, we altered the size of sample 2 (N2) during resampling while maintaining the same population structure. The sc-UniFrac method was observed to be robust with respect to imbalanced dataset sizes, with only minor losses in sensitivity at larger imbalances (detection limit of 5% instead of 2%) **(Figure 2E, F, S2)**. These results demonstrate the robustness of sc-UniFrac, whose performance is independent of chosen parameters.

**Figure 2:**
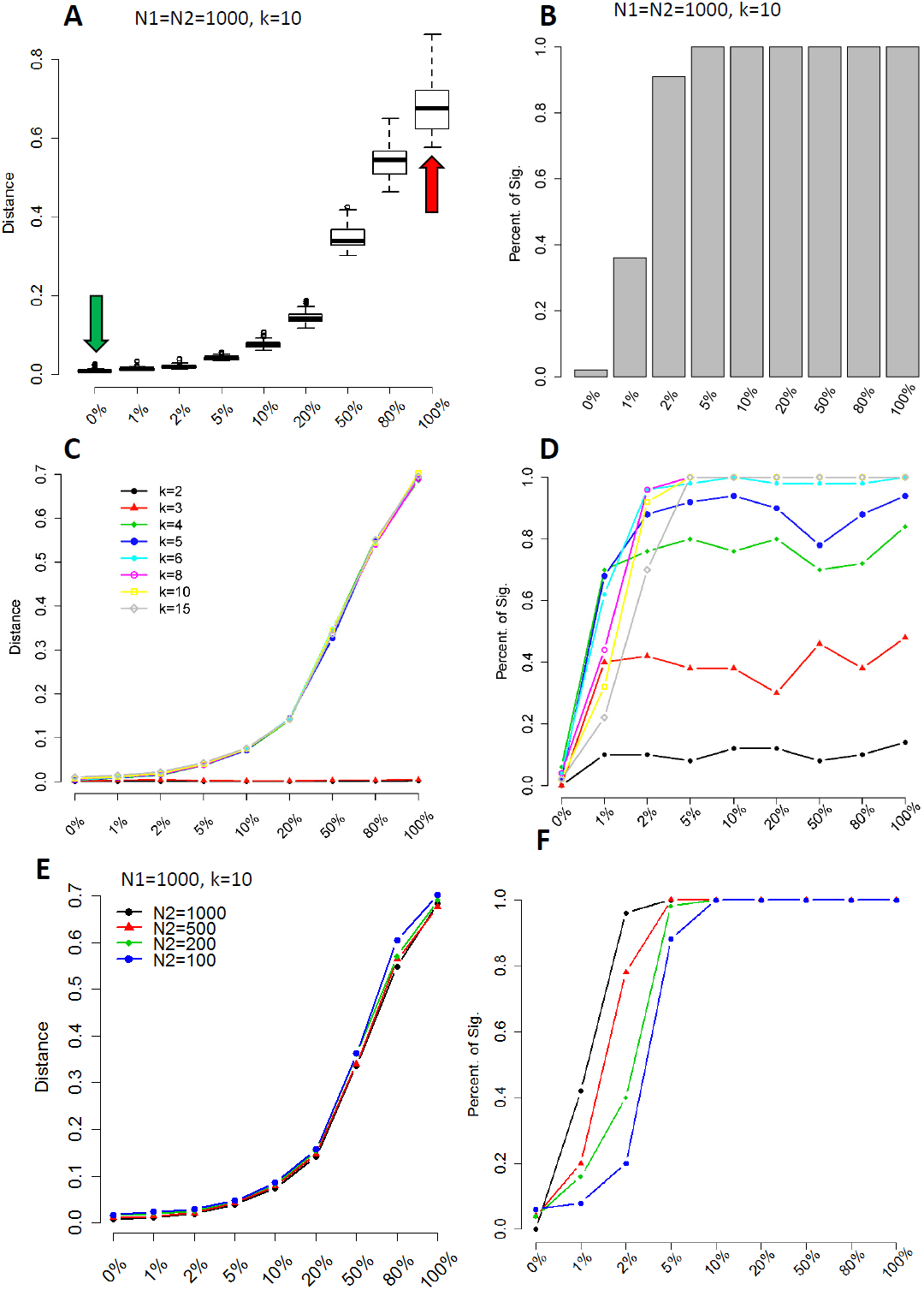
Simulation data reveal sc-UniFrac to be sensitive and robust. (A) Two groups (N1 and N2) of 1000 cells were selected from CD8 and CD4 cells identified in the Wishbone dataset [25]. N1 is always composed of 100% CD8 cells, while N2 is composed of CD8 cells and different proportions of CD4 cells (indicated on X axis). Green and red arrows represent CD8/CD8 (completely similar) and CD8/CD4 (completely dissimilar) comparisons, respectively. Y-axis is the sc-UniFrac distance calculated over n=50 runs with k=10. Boxes represent the first and third quartiles and bars represent maximum and minimum values. (B) Sensitivity of sc-UniFrac evaluated by the fraction of incidences that a statistically significant sc-UniFrac distance was returned over n=50 runs, as a function of increasing dissimilarity between N1 and N2 using the same simulation scheme as A. (C) Mean sc-UniFrac plotted as in A with varying k parameter.(D) Fraction significant sc-UniFrac detected plotted as in B with varying k parameter. (E) Mean sc-UniFrac plotted as in A with N1=1000 but a varying N2 size to determine the effect of dataset size imbalance on sc-UniFrac. (F) Fraction significant sc-UniFrac detected plotted as in B with N1=1000 and varying N2 size.

### Assessment of reproducibility of scRNA-seq data and dissimilarities between different tissue samples

To demonstrate the utility of sc-UniFrac on scRNA-seq data, we generated a series of scRNA-seq datasets with known similarities/differences using inDrop sequencing [1]. These datasets included tissue samples from the mouse colonic epithelium, consisting of both technical and biological replicates, as well as biological replicate samples of the mouse embryonic pancreatic islet collected at E14.5. Three technical replicates (colon1_1, colon1_2, colon1_3) consisted of multiple single-cell fractions collected from one mouse, whose libraries were constructed and sequenced on separate days. Three biological replicates (colon1, colon2, colon3) consisted of samples collected from different mice on different days, but were processed and sequenced together. One sample (colon1) has both technical and biological replicates.

Two traditional strategies to assess the reproducibility of scRNA-seq datasets, for both biological and technical replicates, were performed. One strategy is to compare the median levels of all expressed genes between every sample pair using Spearman correlation analysis. The correlation analysis demonstrated that the technical replicates were more similar amongst each other (mean R= 0.89 +/- SD 0.04), compared with samples amongst biological replicates (mean R= 0.80 +/- SD 0.07) **(Figure S3)**. As expected, the embryonic islets displayed a median gene expression that was the most different when compared to the adult colon (mean R= 0.504 +/- SD 0.07). These results are consistent with expected similarity among different conditions, with technical replicate > biological replicate > outgroup organ. While correlation analysis can quantify the degree of similarity in terms of average transcriptional profile, it provides a very rough estimate and tends to be easily biased towards the dominant population in single-cell landscapes. The other strategy is to do a visual evaluation on how cells from multiple samples are intermixed on a t-SNE plot. Structural differences amongst different samples will be reflected in segregation of data points into separate clusters by sample, while data points from similar samples will appeared together as mixed clusters. Visualization on t-SNE plot showed the same result as median correlation analysis. Technical replicates appeared more intermixed within t-SNE clusters compared with biological replicates **(Figure 3A, B, S4A, B)**. In contrast, pancreatic biological replicates segregate away from samples generated from the colon as both technical and biological replicates **(Figure 3C, D)**. While t-SNE analysis describes the subpopulation structure of the samples, it is not quantitative in that similarities and differences were assessed subjectively by visualization.

**Figure 3:**
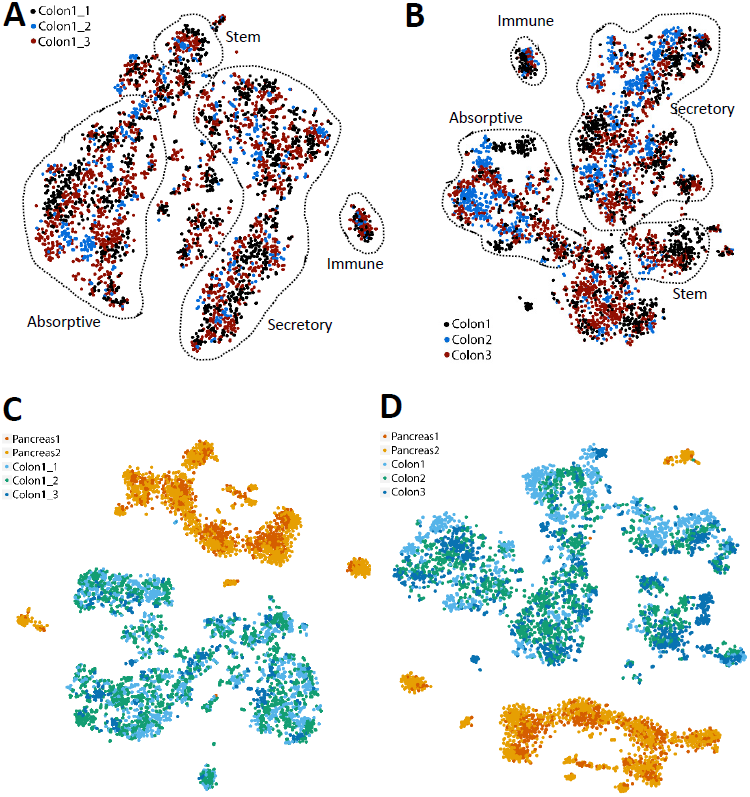
Data landscapes generated from controlled scRNA-seq experiments on technical and biological replicates of different tissues. t-SNE plots of (A) technical and (B) biological replicates of scRNA-seq data generated from the adult murine colonic mucosa. Replicates were combined for t-SNE analyses and labeled with different colors. Outlined populations were identified with canonical markers. t-SNE plots depicting E14.5 pancreatic islet and adult colonic mucosa scRNA-seq data in (C) technical and (D) biological replicates, showing segregation by organ type.

Compared with these two traditional strategies, sc-UniFrac provides an objective, precise, and unbiased metric to quantify compositional dissimilarities across single-cell RNA-seq datasets while taking population structures into account. The calculated sc-UniFrac distance between the colonic and pancreatic datasets was 1, the maximal obtainable distance demonstrating the samples did not share any cell populations **(Figure 4A).** Much smaller, but significant, distances were observed among biological replicates of colonic datasets (**Figure 4A, B,** distance=0.24-0.37), suggesting that they share cell populations but proportional compositional difference can still be detected. Technical replicates appeared the most similar with sc-UniFrac being marginally small without statistical significance (**Figure 4A, B**, distance=0.05~0.09), suggesting that they are comprised of almost identical data points. Importantly, the ordering by similarity across samples was robust to the k parameter **(Figure 4B, C)**. A metric was defined to evaluate the power of sc-UniFrac for discriminating biological replicates from technical replicates (discriminative ability) by subtracting the smallest distance between biological replicates by the largest distance between technical replicates. A positive discriminative ability suggests that sc-UniFrac can discriminate technical from biological replicates. Notably, the k parameter again did not affect the discriminative ability except when k is very small (k<=2) **(Figure 4B, C)**. These results demonstrate the ability of sc-UniFrac to objectively and quantitatively determine dissimilarities between single-cell datasets, as seen by the ordering of samples by their expected similarities (technical replicate > biological replicate > outgroup organ).

**Figure 4:**
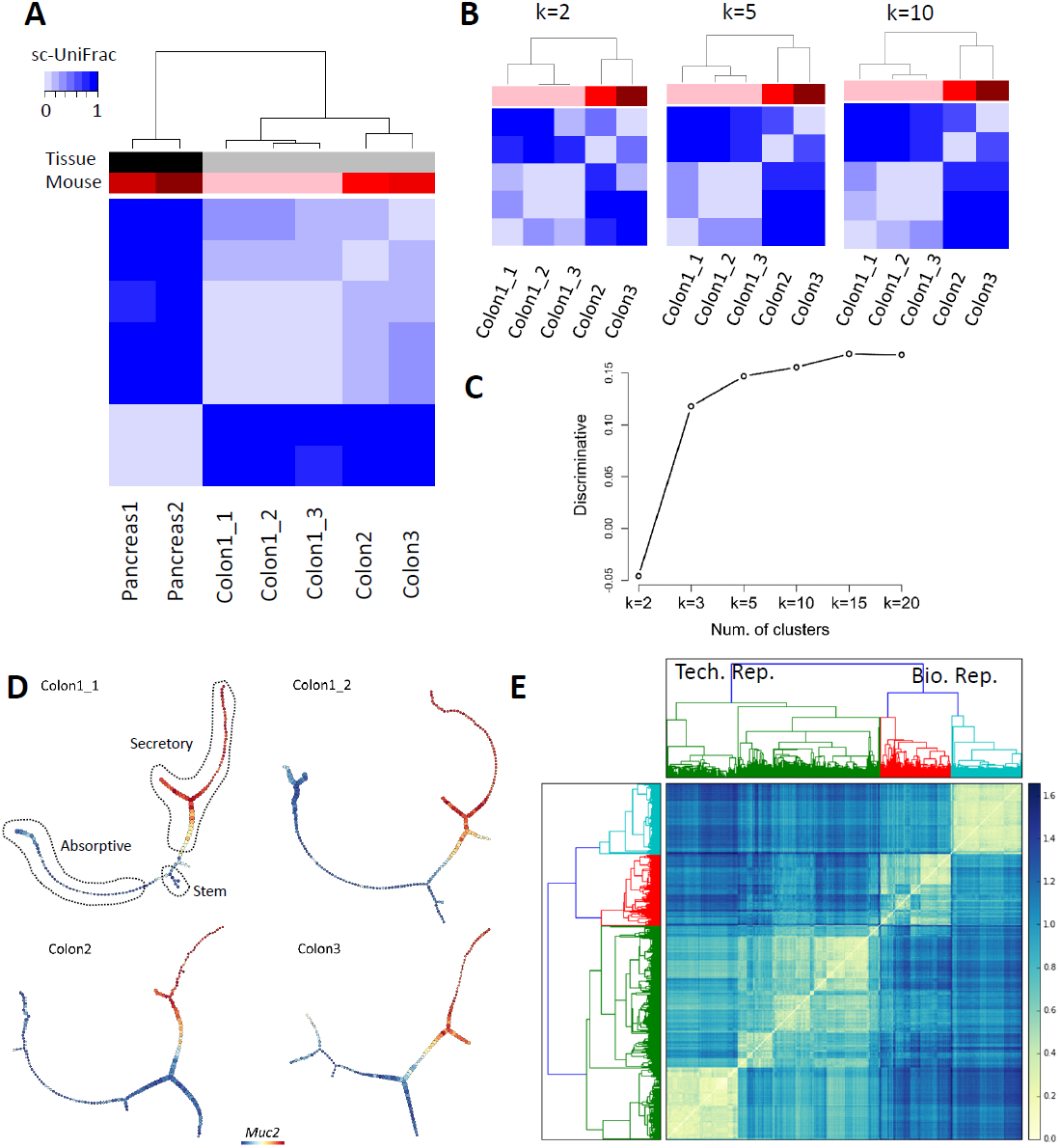
sc-UniFrac can statistically determine dissimilarities between single-cell data landscapes. (A) Hierarchical clustering by sc-UniFrac of scRNA-seq landscapes generated from E14.5 pancreatic islet and adult colonic mucosa (indicated by tissue label), with technical and biological replicates (indicated by mouse label). Heat represents sc-UniFrac distance between two samples. (B) Hierarchical clustering by sc-UniFrac of single-cell landscapes of technical and biological replicates of the colonic mucosa while varying parameter k. (C) Discriminate analysis of sc-UniFrac on biological and technical replicates. Discriminative ability, as defined by the smallest distance between biological replicates minus the largest distance between technical replicates, plotted against k. (D) Representative p-Creode trajectories depicting the colonic epithelial differentiation continuum of 2 technical and 2 biological replicates. Outlined lineages were identified with canonical markers. *Muc2* expression overlay. (E) Hierarchical clustering by p-Creode scoring of trajectories generated from scRNA-seq data of technical (green) and biological (red, cyan) replicates. N=100 resampled p-Creode runs for each dataset were performed and then analyzed together in a single clustering analysis. Heat represents the p-Creode score between two trajectories.

To enable an orthogonal quantitative comparison method to bolster sc-UniFrac results, we used p-Creode trajectory analysis to compare single-cell landscape structures. p-Creode is an algorithm by which a single-cell landscape composed of continuous cell-state data is represented as acyclic graphs to model transition trajectories [19]. The p-Creode score, developed to determine the topological similarity between graphs with differing nodes and edges, can be used to quantify dissimilarities of the trajectory graph outputs generated from different datasets. We revised the p-Creode scoring method to accommodate comparisons of graphs of difference sizes by interpolating between edges connecting nodes, instead of directly matching node positions, between the two test datasets **(Figure S5A-C, and Methods)**. p-Creode was applied to each dataset for 100 times to generate consensus trajectories using data resampling. The modified p-Creode score was used as a distance metric for clustering cell-transitional trajectories created from the resampled datasets. Consistent with sc-UniFrac, p-Creode trajectories amongst technical replicates clustered together using the p-Creode score as a dissimilarity metric, while data from biological replicates were more disparate **(Figure 4D, E)**. As expected, organ specificity drove clustering when pancreatic data were added to the analysis, with all trajectories generated from colonic data clustering together away from pancreatic trajectories **(Figure S5D, S6)**.

p-Creode was designed for data that are distributed as a continuum, and not as distinct clusters. In this regard, comparison can only be made for tissue systems that are transitioning, which is the case for both the adult colonic epithelium and embryonic pancreatic islets compared here. sc-UniFrac does not have this limitation as it can compare between datasets of any distribution, including continuous data as well as discrete populations that are comprised of cells from different lineages. Thus, sc-UniFrac has greater general utility for determining dissimilarities of single-cell datasets in an unsupervised way without prior knowledge of the distribution of the data.

### Identification of cell populations that drive differences between single-cell landscapes within the sc-UniFrac framework

While sc-UniFrac can statistically measure the population diversity between two single-cell landscapes, it also provides an easy and intuitive way to identify the cells that drive the difference. The distance-driving cells are either expanding or contracting populations or even newly emerging populations across conditions. Because sc-UniFrac operates on the statistical basis of shared and unshared weighted branches, one only needs to identify branches that are marked as unshared, characterized by significant proportional shifts of assigned cells between two samples. Unlike the weighted UniFrac algorithm to determine the dissimilarity of microbial landscapes, which identifies the non-zero branches, sc-Unifrac uses a random permutation test to detect those branches whose proportion shift in cell populations cannot happen by chance alone (**Methods**). Illustrating this concept, sc-UniFrac was performed to demonstrate pairwise comparison of scRNA-seq datasets of colonic and pancreatic tissue with k = 10. As expected, technical replicates of the colon with the smallest sc-UniFrac value have mostly shared branches between them, with only one branch with subtle proportional shifts **(Figure 5A)**. In contrast, comparison of the pancreatic and colonic datasets revealed no shared branches, with every unshared branch being highly significant **(Figure 5B)**. Evaluation of unshared branches can easily pinpoint cell groupings that contribute to sc-UniFrac. Here, we focus on group 10, which was entirely contributed by the pancreatic sample. Supervised analysis of differential gene expression revealed the unique gene signatures of these cells compared to colonic populations, which can be identified by canonical marker genes (e.g., group 1 represents deep crypt secretory cells; 2 and 3 are colonocytes; 4 and 5 are goblet cells; 6 are intraepithelial lymphocytes) **(Figure S7)**. Projection of cells from group 10 onto reference cell type gene expression signatures from the Mouse Cell Atlas [3] revealed that individual cells mapped on pancreatic 1) acinar, duct cells, 2) endocrine cells, and 3) immune cells **(Figure 5C)**. These results demonstrate the utility of the branching feature of sc-UniFrac to statistically determine cell populations that drive difference between single-cell landscapes. Notably, none of the methods that we used above for comparison with sc-UniFrac can perform this task.

**Figure 5:**
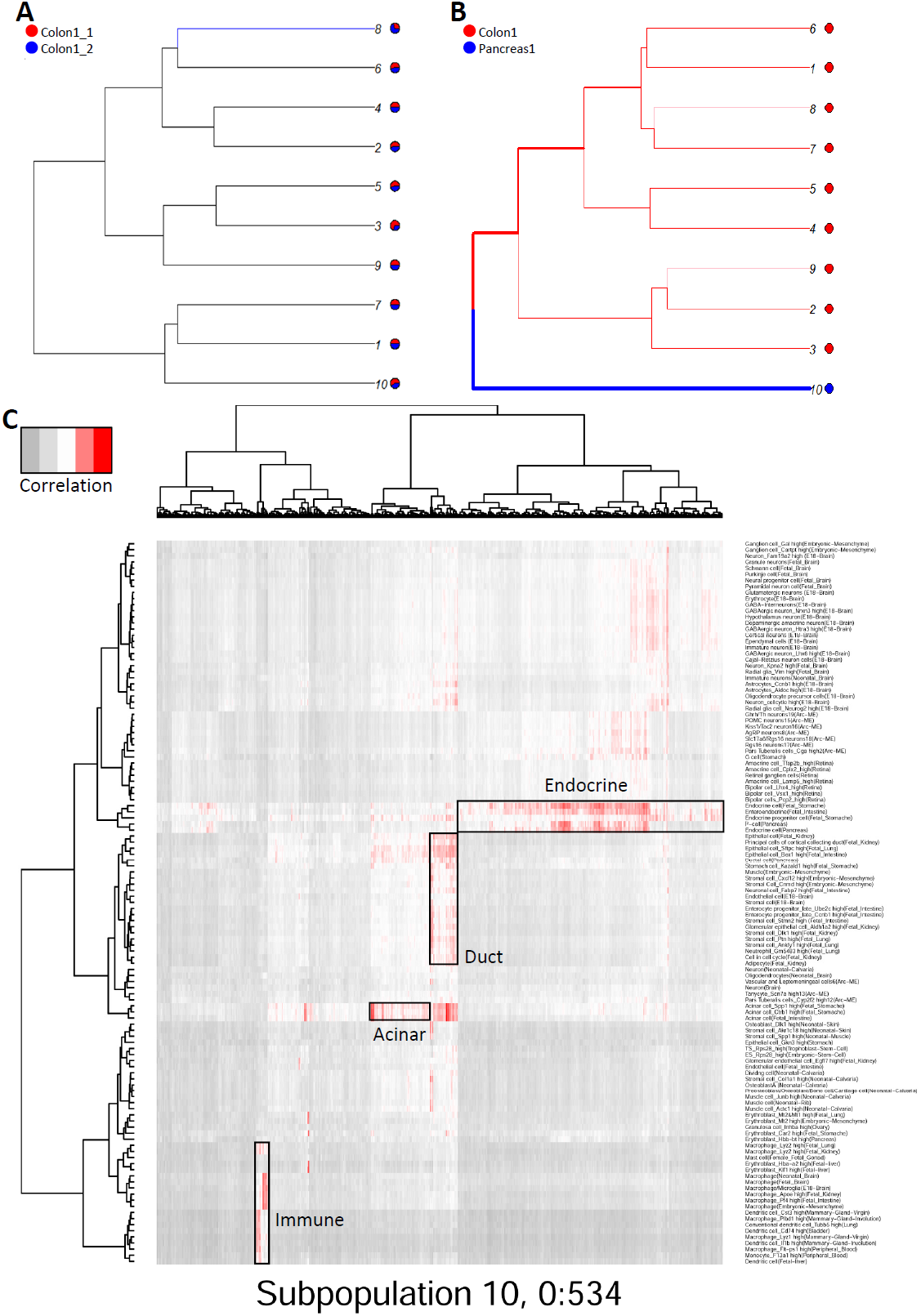
Cells that drive sc-UniFrac can be intuitively identified. (A, B) Branching structure of two single-cell landscapes being scored by sc-UniFrac (k=10), with black representing statistically shared branches, and blue and red representing statistically unshared branches from each of the colored samples. Thickness of branch is proportional to effect size. Comparing between (A) technical replicates and (B) different tissues. (C) Individual cells (columns) from group 10 of B being matched to cell types (rows) referenced from the Mouse Cell Atlas. Heat represents the correlation of gene expression between the cell and the reference using all genes.

### Quantitative evaluation of batch-correction methods by sc-UniFrac

The presence of batch effects is a major and common problem in scRNA-seq experiments, which introduce systematic error and mask underlying biological signals. Removal of batch effects is generally required prior to downstream analysis. Many methods and tools have been developed for batch correction [26–29]. Some methods have been successfully used in bulk RNA-seq [30,31], while other methods were recently developed and specially designed for scRNA-seq [27,32]. While the suitability of batch correction methods may depend on distribution of data that varies from dataset to dataset, the universality of such methods is undefined given that there is no quantitative, objective metric to evaluate batch effect correction in scRNA-seq data. sc-UniFrac, a quantitative measure of cell population diversity in single-cell landscapes, provides a sensitive and objective way to assess the performance of batch correction methods.

We compared three batch removal methods, limma, ComBat and MNN on three scRNA-seq datasets. Limma and ComBat have been widely used for batch correction in bulk experiments, which fit a linear model to determine and then correct the batch effect for each gene. MNN first identifies mutual nearest neighbor pairs between batches and then use these pairs to estimate the batch effect in scRNA-seq data [33]. MNN is expected to perform well when population composition is different across batches. The evaluation of these methods was performed for the following three datasets: 1) HEK293 cells prepared fresh and cryopreserved from two batches [34], 2) our three technical replicates of mouse colonic epithelium, and 3) two separate studies of mouse gastrulation [35,36].

For HEK293 cell line data, a small sc-UniFrac indicating high similarity was observed between freshly isolated and cryopreserved samples within the same batch **(Figure S8)**, indicating minimal technical variation during the cryopreservation process consistent with original findings [34]. In contrast, a large sc-UniFrac was observed between two batches, indicating strong batch effects **(Figure S8)**. All three methods, limma, ComBat and MNN, decreased the sc-UniFrac distance, indicative of batch effect correction **(Figure 6A, S9A-C)**. Among them, limma and ComBat decreased the distances to those approaching to zero, suggesting that batch effects have been completely removed.

**Figure 6:**
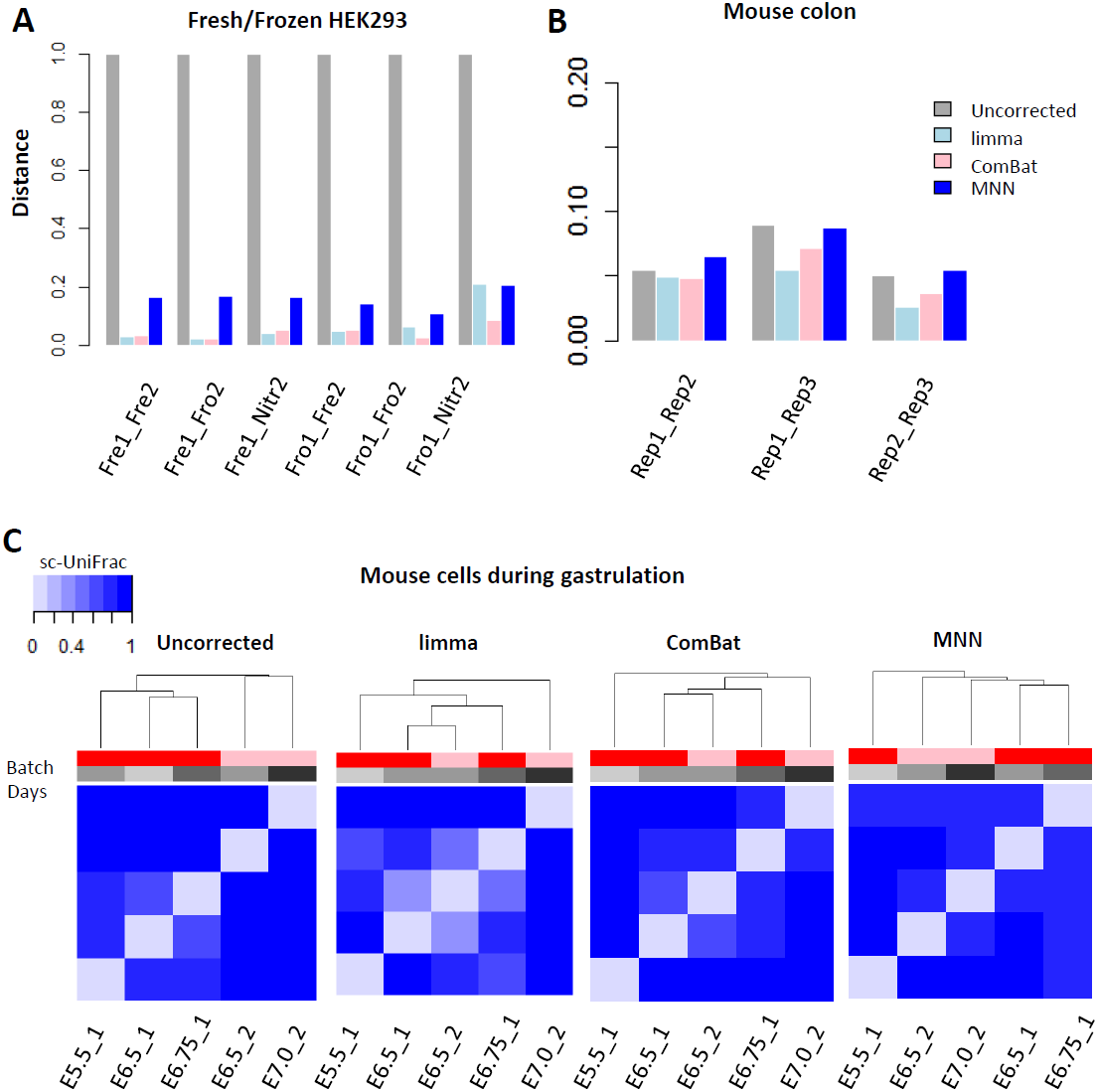
sc-UniFrac can benchmark batch effect removal approaches. (A) sc-UniFrac distance calculated comparing uncorrected and batch-corrected scRNA-seq datasets of HEK293 cells fresh (Fre), frozen at -80 (Fro), or liquid nitrogen flash freezing (Nitr) performed in two different batches [34]. ComBat, limma, and MNN were used for batch correction. (B) sc-UniFrac distance calculated similar to A for technical replicates of the mouse colonic epithelium scRNA-seq data. (C) Hierarchical clustering by sc-UniFrac of uncorrected or batch-corrected scRNA-seq data depicting murine gastrulation from two different studies [37,38]. A gradation of similarity, and hence, clustering, was expected over developmental times from the earliest development stage (E5.5) to the latest stage (E7.5).

In the technical replicates of the mouse colonic epithelium, only very moderate batch effects were observed as indicated by our previous analyses. Due to the initial small differences in batch, limma, ComBat and MNN only moderately removed batch effects further, as seen in the decrease in sc-UniFrac between replicates 1 and 2, and between replicates 1 and 3 **(Figure 6B, S9D-F)**. Batch effects were initially minimal between replicates 2 and 3. In this case, limma and ComBat successfully removed the batch effects (reduced sc-UniFrac to zero), while MNN failed to do but instead introduced additional systematic bias (sc-UniFrac increased) **(Figure 6B)**.

scRNA-seq data of mouse cells during gastrulation were obtained from two studies [37,38], which used plate-based Smart-seq2 and G&T-seq (genome and transcriptome sequencing), respectively. The first study generated scRNA-seq data from mouse embryos at E5.5, E6.5 and E6.75, and the second focused on mouse embryos at E6.5 and E7.0. sc-UniFrac generated on the uncorrected data indicated that the two datasets clustered by studies and not by developmental stages, revealing strong technical variation between the two studies **(Figure 6C, S10A)**. After applying limma, ComBat and MNN, sc-UniFrac generated on the corrected data demonstrated the removal of the technical variation, with cells no longer clustering by studies but by developmental stages. Among the three methods, ComBat achieved the best performance by arranging cells chronologically from the earliest development stage (E5.5) to the latest (E7.5), and achieving the lowest sc-UniFrac between cells from the same stages between the two studies **(Figure 6C)**. This conclusion is supported by t-SNE analysis where E6.5_1 and E.6.5_2 clustered together after limma and Combat, but remained separated after MNN **(Figure S10B-D)**.

From these results, limma and ComBat both outperformed MNN, probably due to identical population composition across batches in all datasets. One of MNN assumptions is that batch effects should be much smaller than biological variation, which may not hold true in these datasets. Additionally, the performance of MNN is dependent on the number of nearest neighbors to consider when identifying mutual nearest neighbor pairs. Choosing the correct parameter would probably improve MNN performance, but this would require prior knowledge of what the correct parameter is. While all methods were able to reduce technical variation, sc-UniFrac was able to quantitatively evaluate the initial batch effects and the performance of batch correction methods.

## Discussion

We developed a new tool, sc-UniFrac, for quantitatively assessing the dissimilarities in cell population structures between two single-cell landscapes. Compared with existing methods, sc-UniFrac has distinct advantages, including: 1) its ability to non-subjectively and quantitatively assess population diversity differences, 2) its precision by taking population structure into account, 3) its statistical rigor based on the available UniFrac framework, 4) its intuitive and statistical robust method to identify disparate cell populations between samples, 5) its flexibility to analyze multiple samples and to add new samples to current analyses, and 6) its ability to handle any dataset with unlimited dimensional representation and any distribution [16]. We have demonstrated the validity of sc-UniFrac using gold-standard datasets where the similarities between datasets are known.

Single-cell technologies provide unprecedented resolution to study heterogeneity in disease, especially in cancer. Intratumor heterogeneity is a key determinant of tumor diagnosis, prognosis, and drug response [39,40]. Although a large amount of effort has been devoted to the genomic [41], transcriptomic [42–44], and proteomic [45] subtyping of cancers in hope for better precision application and/or to better understand the disease, current bulk analyses obscure signals coming from distinct cell populations. Unbiased characterizing cellular diversity in tumor tissues and then using this information to define subtypes provides a unique opportunity to understand cancer. Subtypes based on tumor heterogeneity refine the subtypes defined by the bulk-omics approaches, and may provide additional prognostic and diagnostic value for predicting patient survival and drug response. For current single-cell applications, comparing heterogeneity between multiple samples has been performed manually using t-SNE analysis in conjunction with distinguishing markers to match cell populations across samples qualitatively, which is done in a low throughput fashion with few samples [46]. sc-UniFrac enables quantitative evaluation of cellular diversity among potentially large numbers of samples, which can then be rapidly clustered into different subtypes. Thus, sc-UniFrac can facilitate studies on intratumor and intertumor heterogeneity that reveal the importance of diverse cell populations in tumor progression and drug treatment.

Cell populations will expand, shrink, or emerge as a function of disease subtype, disease progression, or after extrinsic drug perturbations. As a feature of the UniFrac framework, which utilizes a shared branching approach, altered cell populations or states driving compositional difference can be intuitively identified as unbalanced branches in sc-UniFrac. Moreover, difference-driving cells can be further analyzed to identify gene expression signatures, and their identities and behaviors can be inferred based on transcriptomes of previously referenced cell types. Introduction of new cells into the landscape as a result of perturbation, for instance, the infiltration of CD8 cytotoxic T cells into a tumor, can be deciphered by matching (or blasting) [47] difference-driving cells against the transcriptomes of reference cell types [3], as sc-UniFrac has demonstrated here.

There is currently a proliferation of single-cell data analysis tools, where many of them utilize different approaches for achieving the same goal. In response, it is necessary for the single-cell biology community to benchmark the performance of these tools with reference datasets. Batch effect correction is a very important procedure for removing technical variation. The sources of variation can arise from runs on different sequencing lanes, different single-cell encapsulations, ischemic times, or different tissue preparations, even if procedures are performed by the same person. Several tools designed to remove batch effects have been developed specifically for scRNA-seq [20,33,48,49]. A quantitative measure of performance is required for effective benchmarking, and sc-UniFrac now provides a metric where the similarity between single-cell landscapes of a tissue generated from different batches, before and after batch correction, can be evaluated. While evaluation of a tool on any specific datasets can be performed, it should be noted that the performance of any particular tool depends on the assumption of the algorithm and the distribution of the dataset. Thus, different tools may perform better on some datasets than others. sc-UniFrac is a new approach for evaluating single-cell landscape similarity across multiple samples in a quantitative and statistically robust manner. We have demonstrated various applications of this approach and we envision its broad usage as increasing number of scRNA-seq are generated.

## Competing Financial Interests

The authors declare no competing financial interests.

## Acknowledgement

K.S.L is funded by NIDDK (R01DK103831) and NCI (U01CA215798). Q.L., K.S.L. and R.J.C. are funded by P50CA095103 - Vanderbilt GI Special Programs of Research Excellence. C.A.H. is funded by a training grant from NICHD (T32HD007502) and a pre-doctoral F31 from NIGMS (F31GM120940). A.B. and A.J.S. are funded by NIDDK (R01DK103831). R.J.C. is funded by NCI (R35CA197570). Y.S., K.S.L. and R.J.C. are funded by NCI (P30CA068485). G.G. is funded by NIDDK (R01DK065949). The authors would like to thank Dr. Olivia Koues and the VANTAGE core for technical assistance, and members of the Vanderbilt Epithelial Biology Center for helpful discussions.

## Author Contributions

Q.L. developed the algorithm, performed data analysis, supervised the research, and wrote the manuscript. C.A.H. and B.C. developed algorithms, and C.A.H. was involved with writing the manuscript. A.B. and A.J.S. performed scRNA-seq experiments. Q.S. engineered software. J.P. performed scRNA-seq data analysis. G.G. contributed material to the research. R.J.C. and Y.S. intellectually contributed to the research. K.S.L. initiated the development of the method, performed data analysis, wrote the manuscript, and supervised the research.

## Methods

### The sc-UniFrac framework

sc-Unifrac is freely available as an R package at https://github.com/liuqivandy/scUnifrac. sc-Unifrac includes four main steps: 1) data processing: scRNA-seq data were first normalized by library size per cell (total number of UMIs) and log-transformed; 2) tree construction: highly variable genes were selected (user defined: default 500). Dimension reduction was performed using PCA (user defined: default 4 PCs). A hierarchical tree representing cell population structure was built by clustering via average linkage, and the upper portion of the tree was defined by cutting off the connections at k clusters (user defined: default k=10); 3) quantification of cell population diversity: the sc-UniFrac distance was calculated by weighted branch sharing, and statistical significance was assessed by permutation testing; 4) identifying populations that drive sc-UniFrac by querying the shared branching structure: gene expression signatures were derived for matching against reference cell type signatures. sc-Unifrac generates a report to summarize the results, including the sc-UniFrac distance on population diversity, statistical significance, cell population structures, gene expression signatures in each altered population, and their match to reference cell types **(Supplementary File)**.

### sc-UniFrac distance calculation

sc-UniFrac (*D*) is calculated as

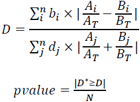

Here, *n* is the total number of branches in the tree. *b*_*i*_ is the length of branch i. *A*_*i*_ and *B*_*i*_ are the number of cells than descend from branch *i* in the two samples A and B, respectively. *A*_*T*_ and *B*_*T*_ are the total number of cells in two samples A and B, respectively. 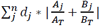 is the average distance of each cell from the root, used to normalized the distance from 0 to 1. *D\** is the distance based on permuted data, while *D* is the observed distance. *N* is the total number of permutations.

Unshared branches are occupied by populations with statistically significant shifts between samples. The proportion shift of a cell population *i* is defined as 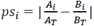 with statistical significance achieved if *Ps*_*i*_ > *Ps\**, where *Ps\** is the proportion shift in permuted datasets.

### Signature matching to reference cell types

The signature composed of under- or overexpressed genes associated with a cell population was defined by limma comparisons with other populations. For predicting cell types, each cell of a given cluster was matched to the 894 cell type references from the Mouse Cell Atlas [3]. Matching was performed by deriving a Pearson correlation of all genes between the query cell and the reference cell type transcriptome.

### Modified scoring of p-Creode trajectories

The p-Creode score was originally designed to compare p-Creode trajectories as a means to evaluate robustness of the calculated trajectories and also to arrive at a representative trajectory over multiple bootstrapped runs on the same single-cell data set. Thus, it was developed to assess the dissimilarity between trajectories of largely the same sizes. Here, we use p-Creode scoring to compare different datasets, which can generate trajectories of different sizes. To make the p-Creode score size-invariant, we modified how nodes are transformed from one graph to another, while maintaining the scoring approach outlined in [19]. Previously, when a node was not contained in both graphs, a node transformation was performed by translating the node in the second graph to the closest node in the first graph with a penalty **(Figure S5A)**. To eliminate excess penalty when a dense graph is transformed into a sparse graph and vice versa **(Figure S5C)**, the scoring routine was updated to allow for transformations into the edges as well as nodes in the reference first graph. More formally, the transformation penalty is the minimum distance to the closest edge projection between two nodes in opposing graphs plus the remaining graph edge distance to the closest node along the path of the pairwise comparison **(Figure S5B)**.

### Batch-Correction methods

We compared three batch-correction methods, limma, ComBat and MNN. For limma, we used removeBatchEffect function in the limma package, which fits a linear model to the data and then removes the component due to the batch effects [31]. For ComBat, we used the ComBat function in the sva package, which uses Empirical Bayes methods to adjust for both the mean and variance differences across the batches [30]. MNN identifies the mutual nearest neighbors between batches and uses them to estimate and remove the batch effect [33]. We performed the mnnCorrect function in the scran package. We set the number of nearest neighbors to consider to be 20.

### Single cell RNA-seq data sources

We used four single-cell datasets in this study. 1) Wishbone dataset [25]. This is a mass cytometry dataset characterizing the mouse thymus during T cell development, where lymphoid progenitors differentiate to either CD8^+^ or CD4^+^ cells. Data on about 250,000 cells on 37 surface markers and transcription factors were generated. We removed the DN cell population and included only CD8^+^ or CD4^+^ cells labeled by Wishbone. We then simulated population mixtures by randomly sampling CD8^+^ and CD4^+^ cells. 2) Fresh and frozen HEK293 scRNA-seq datasets [34]. Single cell transcriptomes of the fresh and cryopreserved HEK293 cells were generated by MARS-seq, which included about 50 cells in each sample. The UMI-filtered read counts were downloaded from GEO with accession number GSE85534. 3) Technical and Biological replicates from mouse colon and pancreas tissues. scRNA-seq of colonic and pancreatic tissues were generated by inDrop platform, with datasets sizes ranging from 463-1652 cells in colonic and 534-1099 cells in pancreatic tissues (Methods described below). The UMI-filtered read counts and raw data are available from GEO with accession numbers GSE102698, GSE114044. 4) Mouse gastrulation [37,38]. scRNA-seq data of mouse cells during gastrulation were obtained from two studies. Mohammed et al. isolated single cells from mouse embryos at different stages and generated scRNa-seq data using G&T-seq. We selected data from the E5.5 (267 cells), E6.5 (168 cells) and E6.75 stages (82 cells). Count tables were downloaded from GEO with accession number GSE100597. Scialdone et al. used Smart-seq2 to profile 1,205 cells of gastrulating mouse embryos. We selected data from the E6.5 (502 cells) and E7.0 stages(138cells).Gene counts were downloaded from http://gastrulation.stemcells.cam.ac.uk/scialdone2016. Ensembl gene IDs were mapped to mouse gene symbols using the biomaRt package.

### Mouse Experiments

Animal experiments were performed under protocols approved by the Vanderbilt University Animal Care and Use Committee and in accordance with NIH guidelines. Wild type mice (C57BL/6) were euthanized in an approved fashion prior to dissection and tissue harvesting.

### Single-cell RNA sequencing of colonic and pancreatic tissues

Single cell suspensions of colonic epithelium were prepared by chelating (3mM EDTA, 1mM DTT) distal colon segments at 4°C for 45 minutes followed by shaking off crypts [19,50]. Isolated crypts were then dissociated into single cells using a DNAse1/collagenase enzymatic cocktail (2.5mg/ml DNAse1, 2mg/ml collagenase) at 37°C for 20 minutes. Crypt fragments were further dissociated into single cells using a 27½ gauge needle. Cell suspensions were washed 2x with cold PBS to remove debris and enriched for live cells using a Miltenyi MACS dead cell removal kit. Live cell concentration was counted based on Trypan Blue positive cells and a solution of 150,000 cells/ml was prepared for encapsulation. To maintain live cell viability, 18ul of Optiprep was added per 100ul of cell solution prior to encapsulation.

Dissociation of pancreatic buds from E14.5 control embryos was performed using previously published protocols [51]. Briefly, pancreatic buds were dissected from control embryos and trypsinized followed by MACS sorting. Single cell suspensions from multiple embryonic buds were prepared and a cell solution of 20,000 cells was prepared for encapsulation.

Single cell encapsulation was performed using the inDrop platform (1CellBio) with an *in vitro* transcription library preparation protocol, as previously described (cite Klein et al 2015). inDrop utilizes CEL-Seq in preparation for sequencing and is summarized as follows: 1) Reverse transcription (RT), 2) ExoI nuclease digestion, 3) SPRI purification (SPRIP), 4) Single strand synthesis, 5) SPRIP, 6) T7 *in vitro* transcription linear amplification, 7) SPRIP, 8) RNA fragmentation, 9) SPRIP, 10) primer ligation, 11) RT, and 12) library enrichment PCR. Number of cells encapsulated was calculated by approximating the density of single cell suspension multiplied by bead loading efficiency during the duration of encapsulation. Each sample was estimated to contain approximately 2500 encapsulated cells.

Following library preparation, as described above, the samples were sequenced using Nextseq 500 (Illumina) using a 150bp paired-end sequencing kit in a customized sequencing run [19]. After sequencing, reads were filtered, sorted by their designated barcode, and aligned to the reference transcriptome using InDrops pipeline. Mapped reads were quantified into UMI-filtered counts per gene and barcodes that correspond to cells were retrieved based on previously established methods [1]. From approximately 2500 cells encapsulated, ~1800 cells were retrieved per sample.

### Experimental Design

Technical replicates were different single-cell encapsulations collected from the same mouse colon, but prepared and sequenced on different days. Biological replicates were tissues collected from different mice on different days but sequenced in the same run.

## Supplementary Figure Legends

**Supplementary Figure S1:**
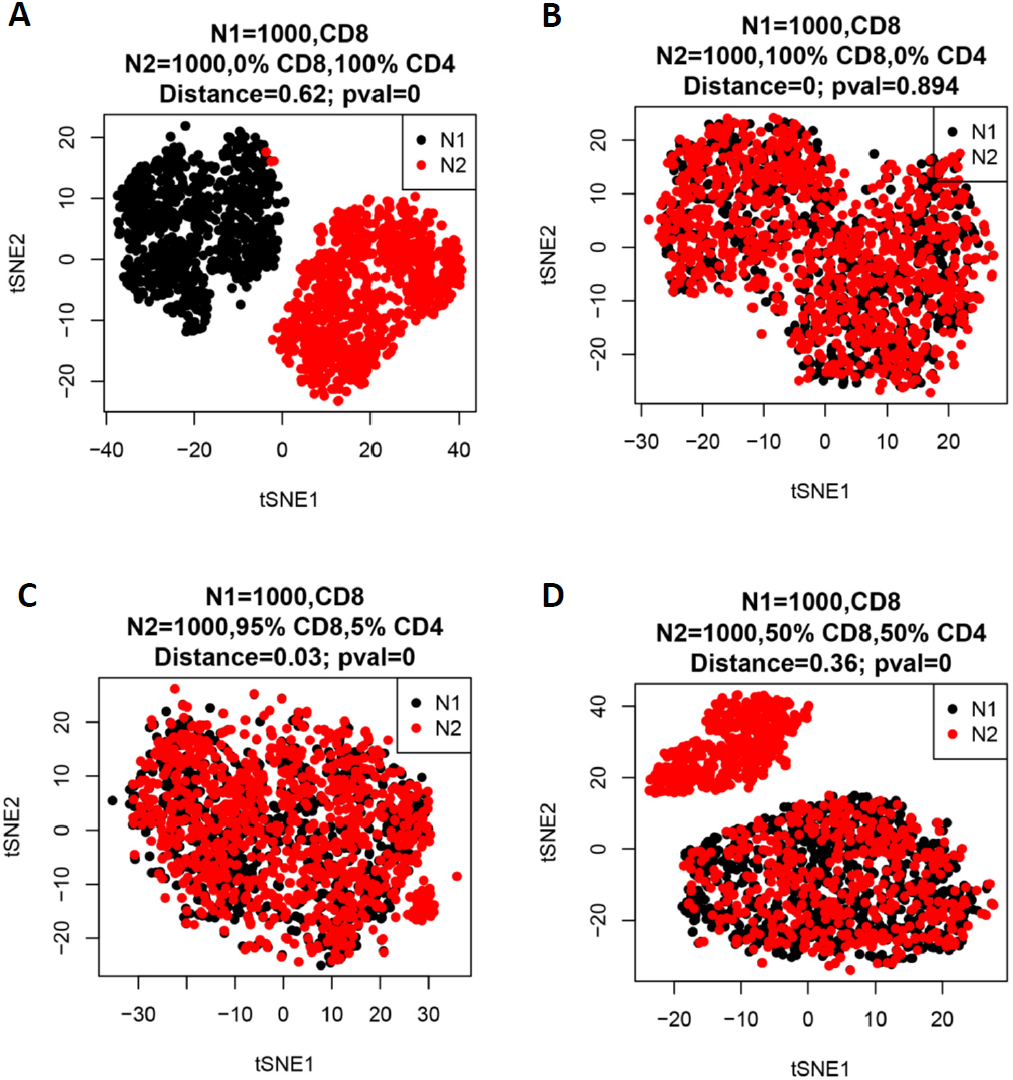
t-SNE plot of two 1000-cell populations simulated by resampling from CD4 and CD8 cells. [25]. One cell population (N1) included only CD8 cells, while the other cell population N2 was comprised of proportional mixtures of CD4 and CD8 cells. (A) N2 = 100% CD4 cells. (B) N2=100% CD8 cells. (C) N2=95% CD8 cells; 5% CD4 cells. (D) N2=50% CD8 cells; 50% CD4 cells.

**Supplementary Figure S2:**
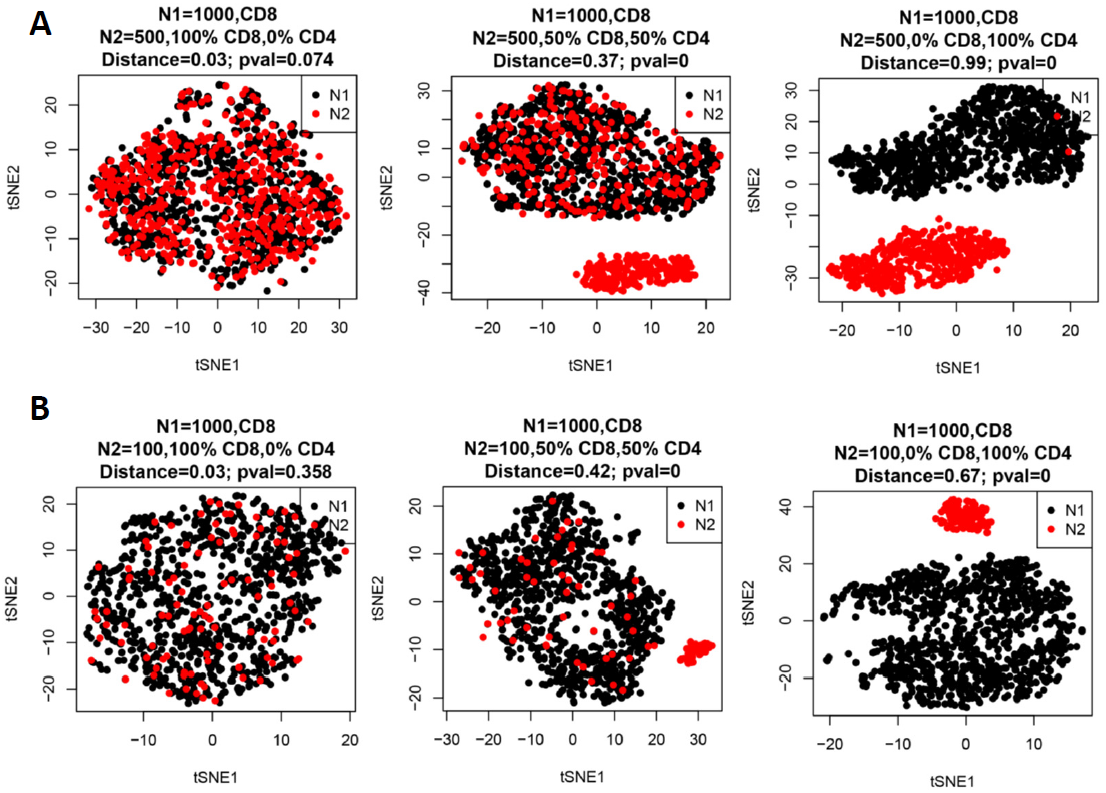
Distribution of data when dataset size is imbalanced. t-SNE plots of similar simulations as Figure S1 with N2 being 100% CD8 cells, 50/50 CD8/CD4 cells, and 100% CD4 cells, going from left to right. Altering the size of N2 to be (A) 500 and (B) 100. N1 remains at 1000 cells.

**Supplementary Figure S3:**
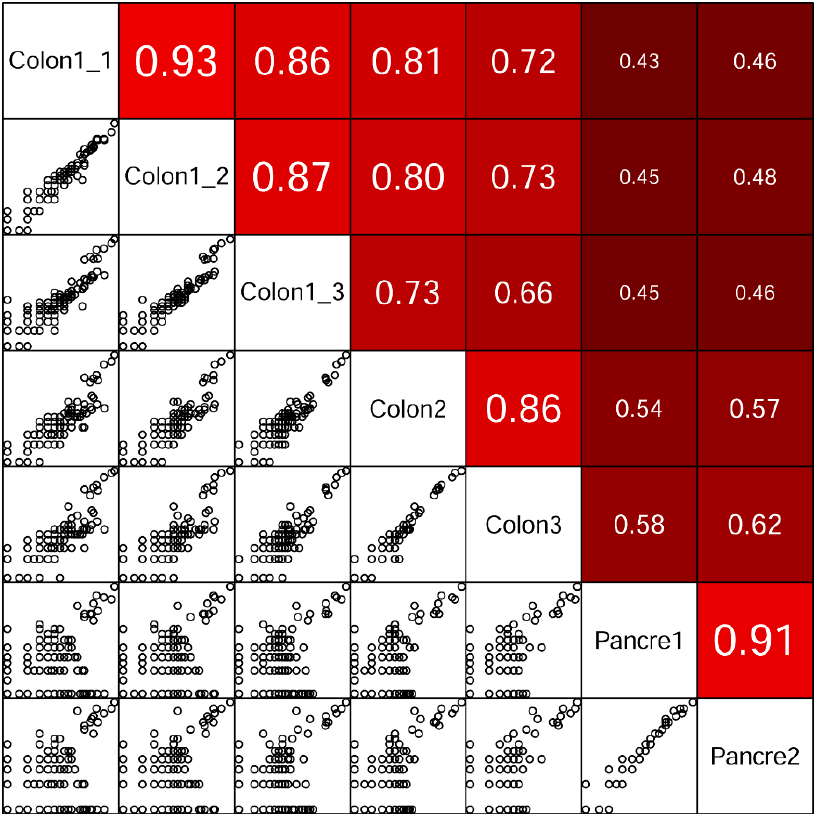
Bulk analysis to demonstrate the ordering of similarity between scRNA-seq data from technical and biological replicates of the colon versus the pancreatic islet. Gene correlation analysis where scRNA-seq data were averaged to generate bulk values. Each data point (on the lower triangle plots) represents a gene whose log expression level was plotted between the two samples being compared. Upper triangle plots are calculated correlation coefficients.

**Supplementary Figure S4:**
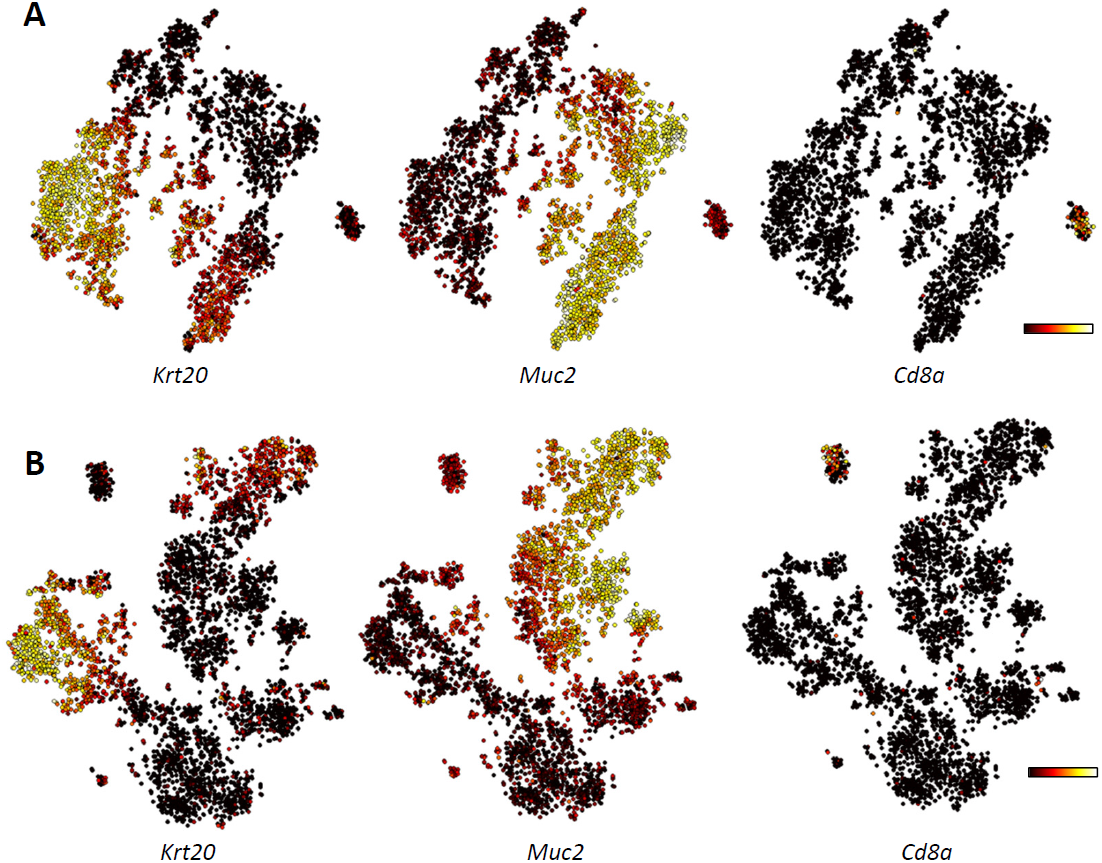
t-SNE analysis of scRNA-seq data from technical and biological replicates of the colon, and the pancreatic islet. *Krt20* depicting the absorptive lineage, *Muc2* depicting the secretory lineage, and *Cd8a* depicting immune cells overlaid on t-SNE plots of scRNA-seq data generated from the adult murine colonic mucosa with (A) technical and (B) biological replicates.

**Supplementary Figure S5:**
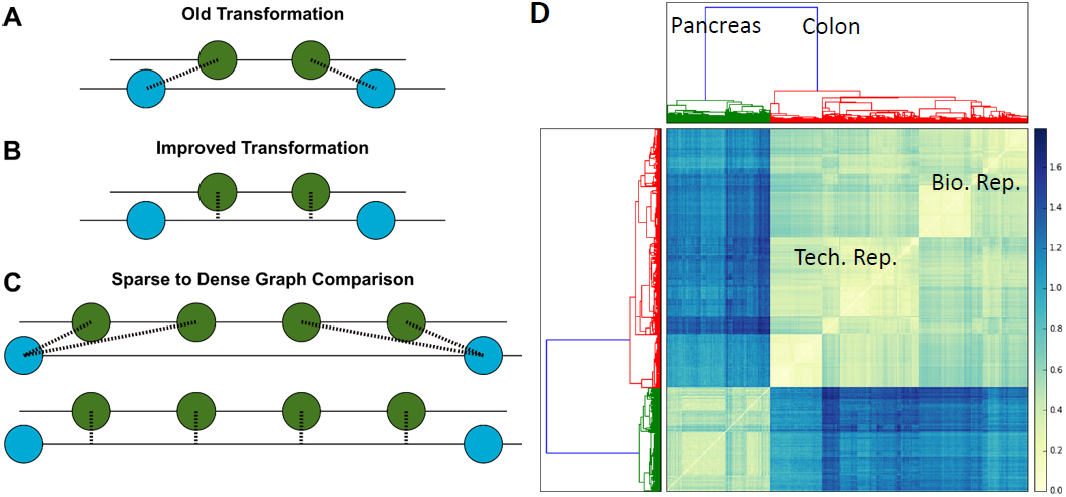
p-Creode scoring can quantify differences between trajectories constructed from continuous single-cell data. (A) Scheme of old node-to-node projection strategy used for the previous p-Creode scoring approach [19]. Dotted line represents Euclidean distance penalty of each transformation. Green and red nodes are from different trajectories. (B) Scheme of new node-to-edge projection strategy used for the current p-Creode scoring approach. (C) Demonstration of excess penalization using the previous p-Creode scoring strategy when there is an imbalance in dataset size resulting in different numbers of nodes in the trajectory (top) versus more realistic penalization with the current approach (bottom). (D) Hierarchical clustering by p-Creode scoring of trajectories generated from scRNA-seq data of E14.5 pancreatic islet (green – biological replicates) and adult colonic mucosa (red – technical and biological replicates). N=100 resampled p-Creode runs for each dataset were performed and then analyzed together in a single clustering analysis. Heat represents the p-Creode score between two trajectories.

**Supplementary Figure S6:**
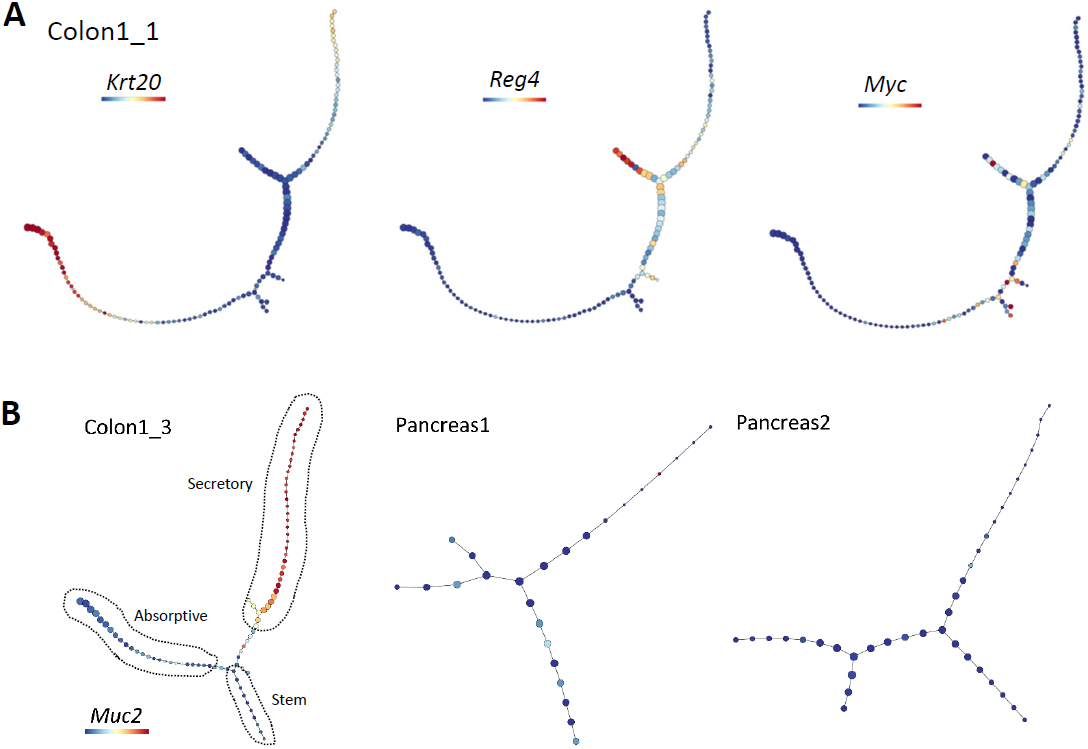
p-Creode trajectory analysis of scRNA-seq data from technical and biological replicates of the colon, and the pancreatic islet. (A) *Krt20* depicting colonocytes, *Reg4* depicting deep crypt secretory cells, and *Myc* depicting stem and progenitor cells overlaid on a representative p-Creode trajectory of scRNA-seq data generated from the murine colonic epithelium. (B) Representative p-Creode trajectories depicting colonic and pancreatic islet differentiation. Outlined lineages were identified with canonical markers. Overlay of *Muc2* transcript level, which was not expressed in the pancreatic islet.

**Supplementary Figure S7:**
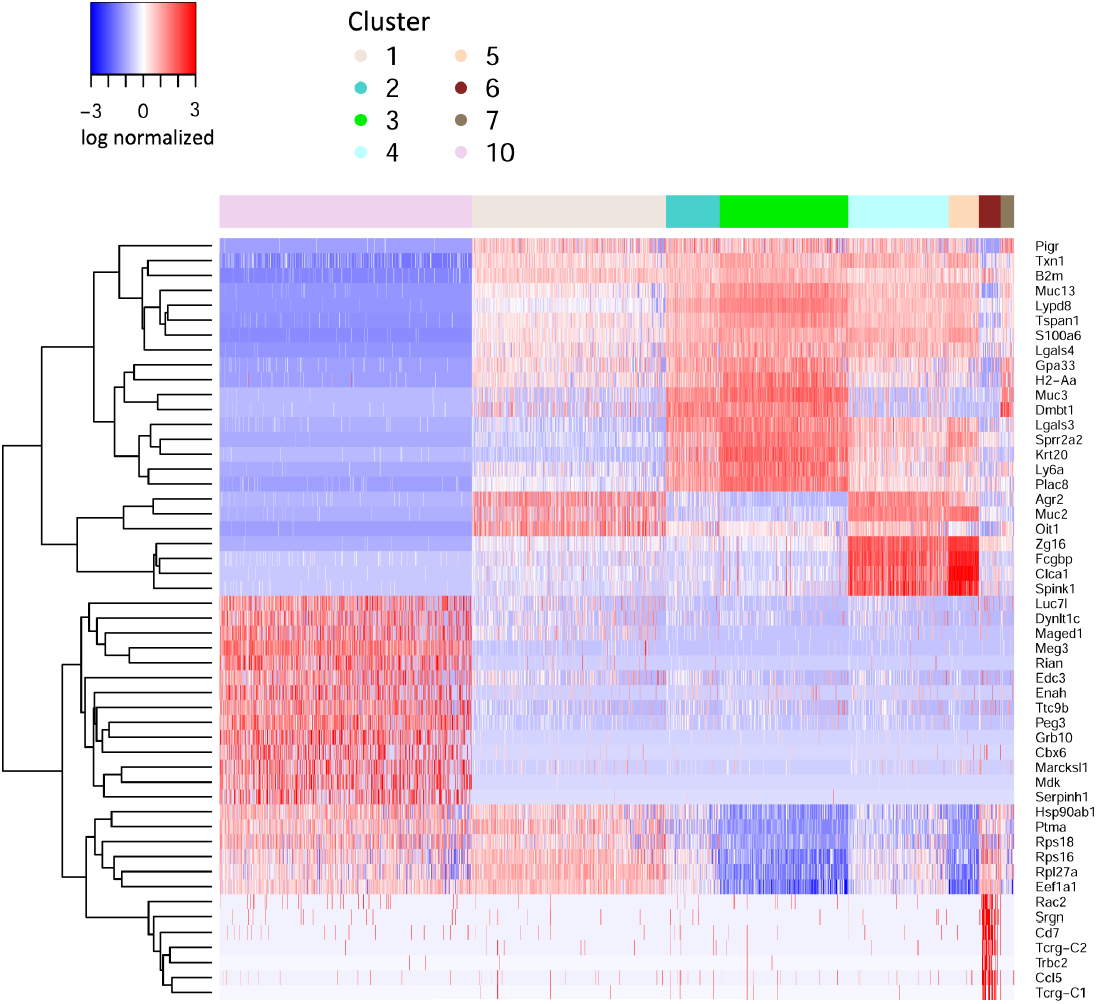
Gene signature extraction from different cell groups. Differential expressed gene identified by limma for each of the 10 groups in Figure 5.

**Supplementary Figure S8:**
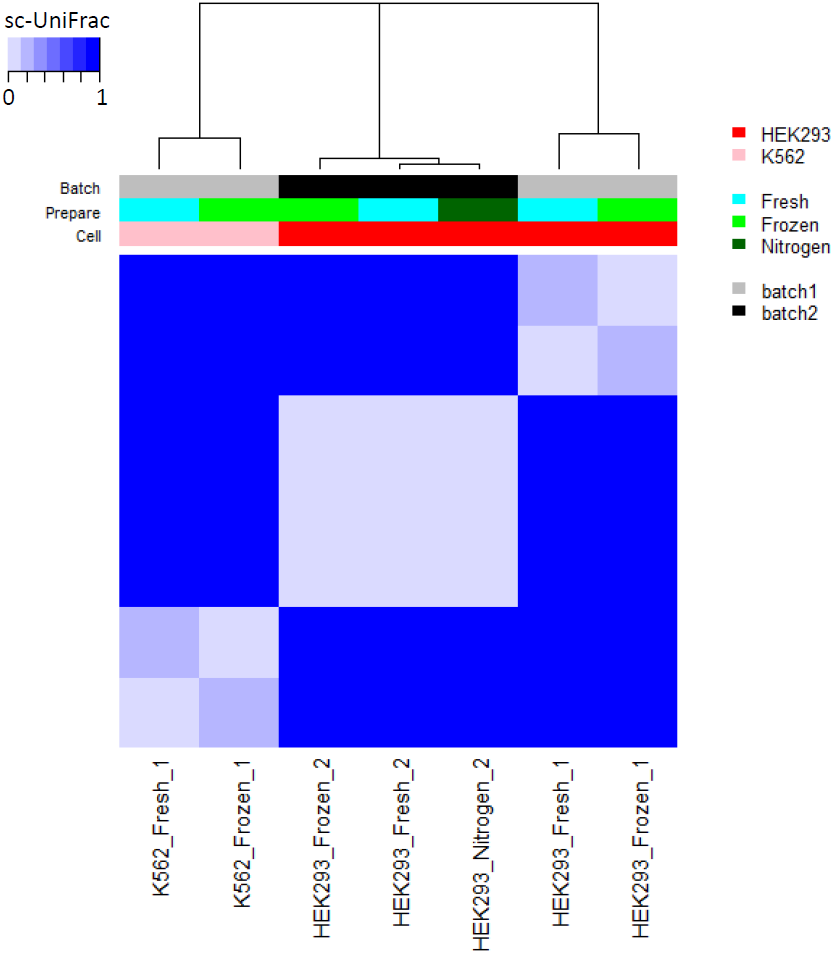
Comparing scRNA-seq data from frozen or freshly prepared samples from different batches. [34]. Hierarchical clustering by sc-UniFrac of scRNA-seq data from cell lines that are prepared differently. Heat depicts the sc-UniFrac distance between 2 samples. The results are consistent with the original study, which shows that the freezing process did not alter transcriptional profiles. In contrast, batch effects have a larger impact on the transcription profiles than the freezing process.

**Supplementary Figure S9:**
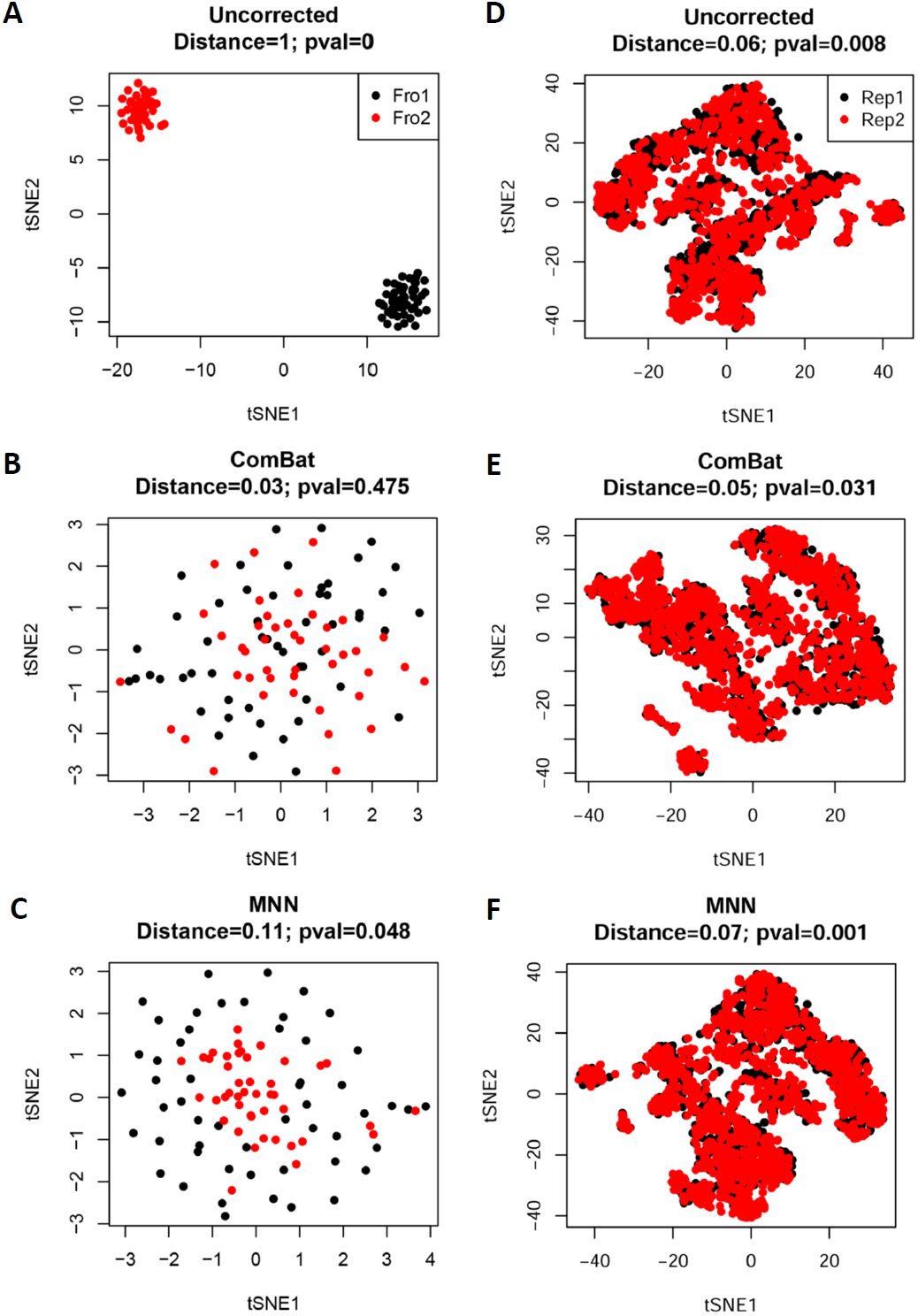
The effects of batch correction. t-SNE analysis of scRNA-seq data from cell lines prepared from two batches (Frozen 1 and 2) [34] (A) uncorrected, (B) corrected by ComBat, and (C) corrected by MNN. t-SNE analysis of scRNA-seq data from the colonic mucosa from two technical replicates (Replicates 1 and 2). sc-UniFrac distance between the samples and p-value noted.

**Supplementary Figure S10:**
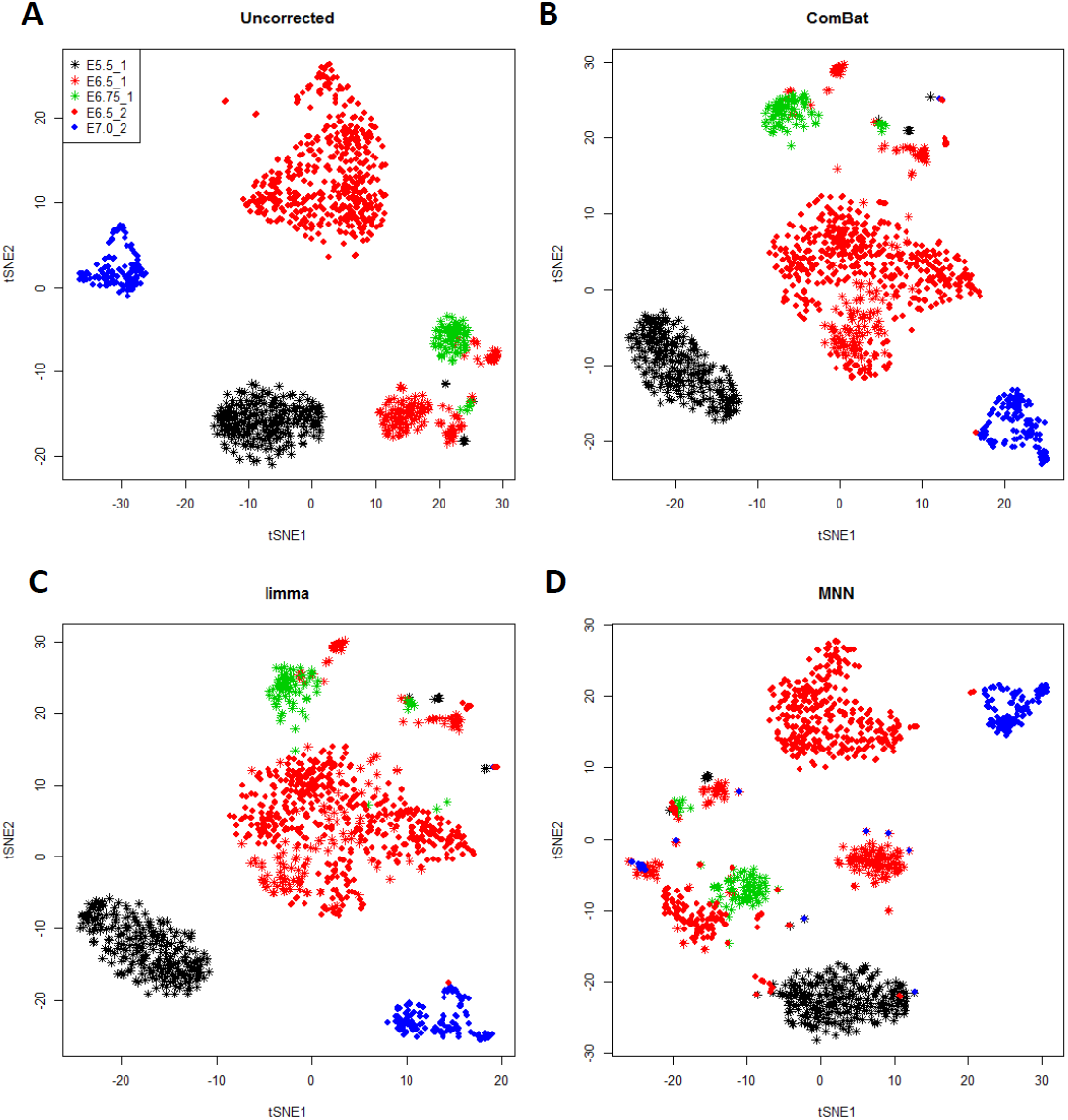
Batch correction applied on data from different studies to align samples according to developmental time. t-SNE analysis of scRNA-seq data depicting mouse gastrulation, with colors representing developmental time and shapes of data points representing the two studies [37,38]. For instance, all red data points should cluster together. Analysis performed on (A) uncorrected, (B) ComBat corrected, (C) limma corrected, and (D) MNN corrected data.

## 1 Overview

**scUnifrac is to quantify cell subpopulation diversity between two single-cell transcriptome profiles**, which calculates the distance, estimates the statistical significance of the distance, identifies the subpopulations that are significant different between two profiles, finds genes that mark subpopulations, and predicts cell types of subpopulations. If two single-cell RNA-seq profiles have identical cell subpopulation structures (the same subpopulations with the same proportion), the distance is zero, whereas, the distance is one if completely different subpopulations.

- ***Results Summary*** - The distance, the statistical significance (p-value), the subpopulations that are different in two profiles, and phylogenetic tree plot and tSNE plot to show the subpopulation structures.
- ***Subpopulation view*** - heatmap of gene expression that mark the statistical significant subpopulations, and predicted cell types of these subpopulations based on Mouse Cell Atlas [Han et al., 2018, Cell, 1091-1107] if scRNA-seq are derived from mouse samples.

## 2 Results Summary

Subpopulation structure:

**Table 1.** The number of cells in each subpopulation in two scRNA-seq profiles.

| Subpopulation | 1 | 2 | 3 | 4 | 5 | 6 | 7 | 8 | 9 | 10 |
| --- | --- | --- | --- | --- | --- | --- | --- | --- | --- | --- |
| Colon | 411 | 113 | 272 | 213 | 64 | 46 | 28 | 7 | 8 | 0 |
| Pan | 0 | 0 | 0 | 0 | 0 | 0 | 0 | 0 | 0 | 534 |

**Table 2.** Significantly different subpopulations and their proportion difference between two scRNA-seq profile.

| Subpopulation | 10 | 1 | 3 | 4 | 2 | 5 | 6 | 7 |
| --- | --- | --- | --- | --- | --- | --- | --- | --- |
| Proportion difference | 1.00 | 0.35 | 0.23 | 0.18 | 0.10 | 0.06 | 0.04 | 0.02 |

**Figure 1:**
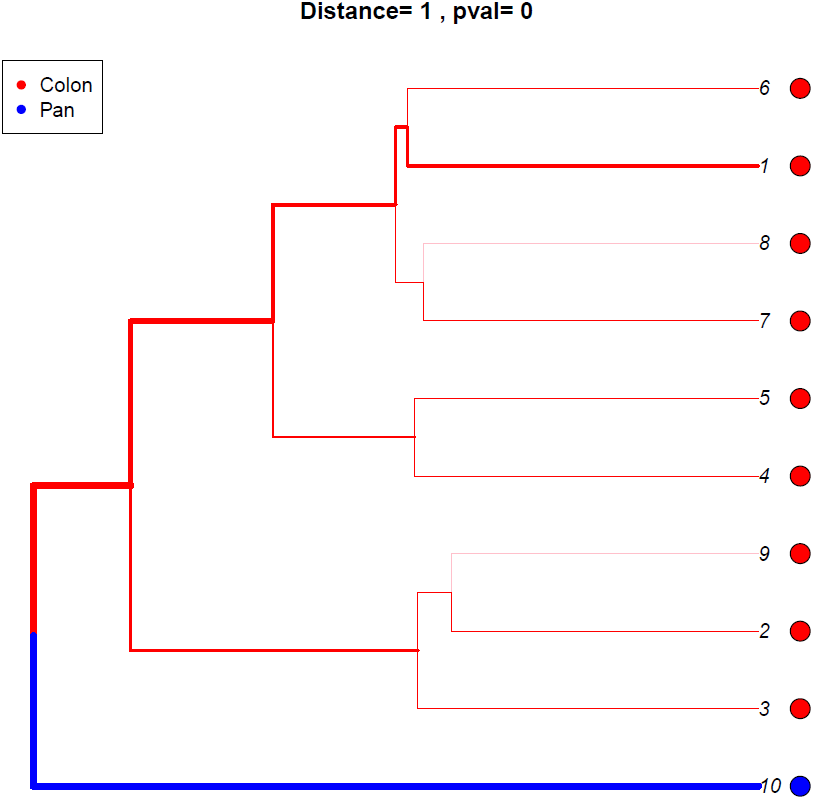
Phylogenetic tree plot of subpopulation structures. Cell subpopulations that are enriched in the first scRNA-seq profile labeled as red, while subpopulations enriched in the second profile are labeled as blue. The line width is proportion to the enrichment. Thicker lines suggest more enriched in one profile. Cell subpopulations that are similar in both samples are labeled as black.

**Figure 2:**
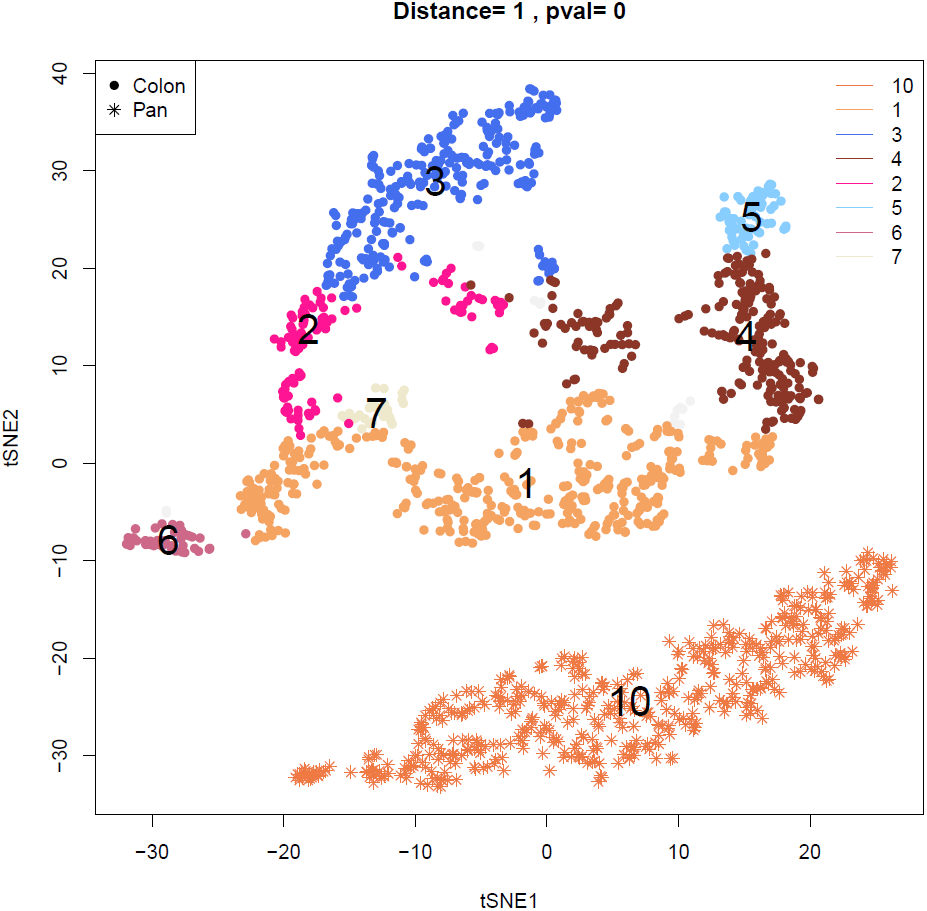
tSNE plot of subpopulation structures. Each subpopulation that are different between two profiles are denoted as different colors, and subpopulations that are similar between two profiles are denotes as gray. Cells from the first profile are denoted by circle, while cell from the second profile are denoted by star.

## 3 Subpopulation View

### 3.1 The cell subpopulation 10

**Figure 3:**
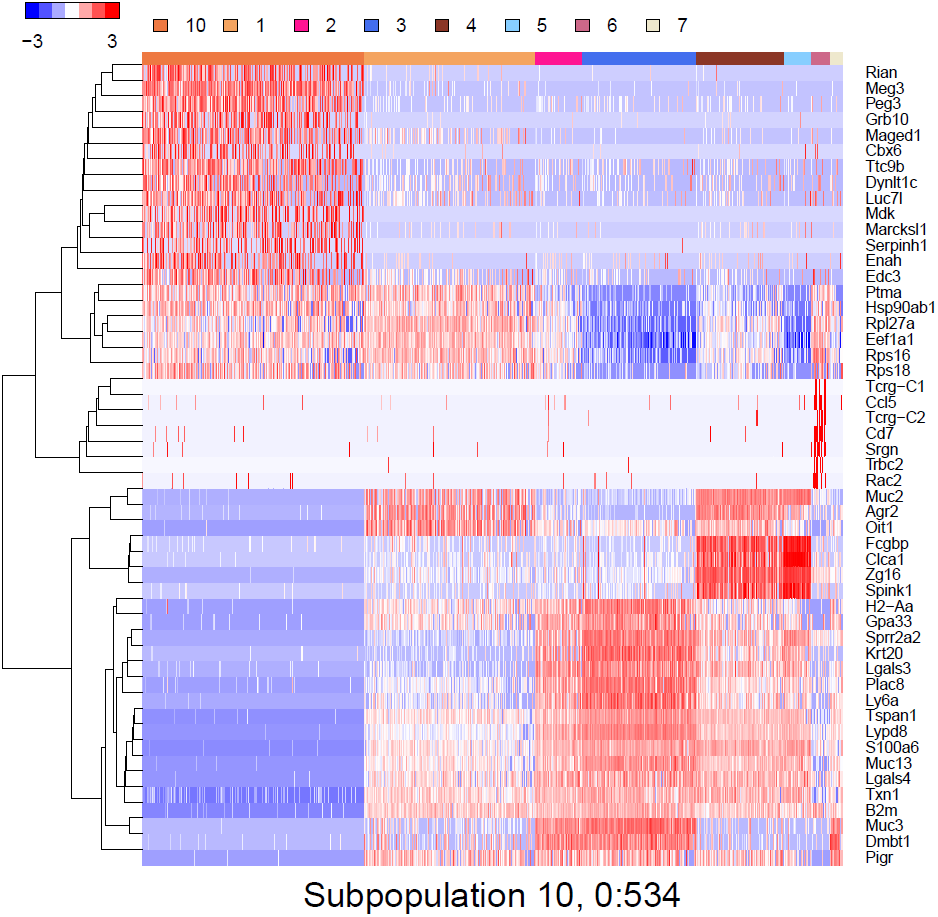
Heatmap of gene expressions that mark the cell subpopulation 10

**Figure 4:**
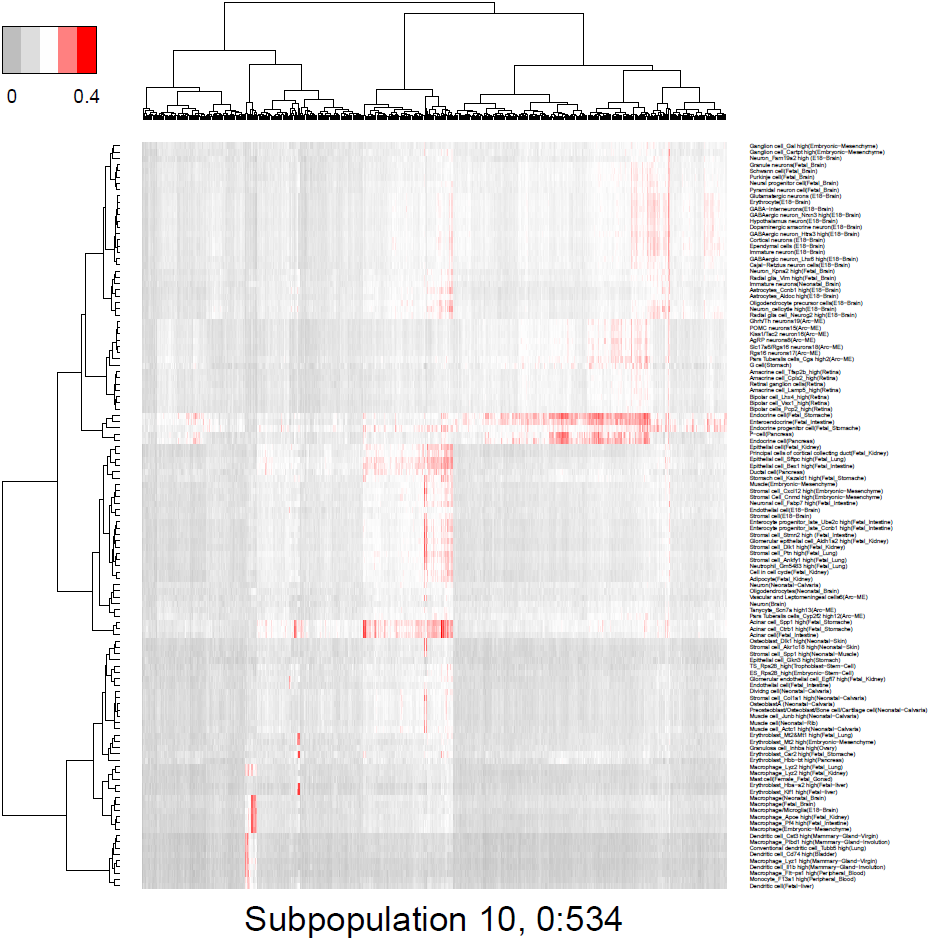
Predicted cell types of the cell subpopulation 10

### 3.2 The cell subpopulation

**Figure 5:**
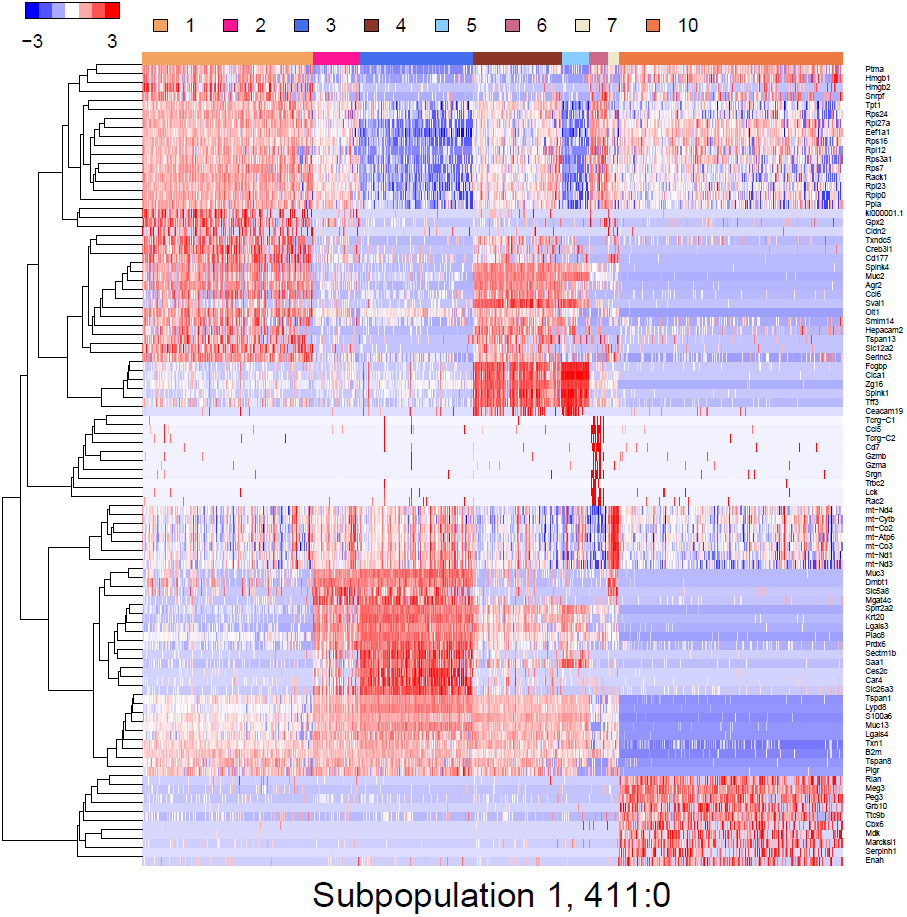
Heatmap of gene expressions that mark the cell subpopulation 1

**Figure 6:**
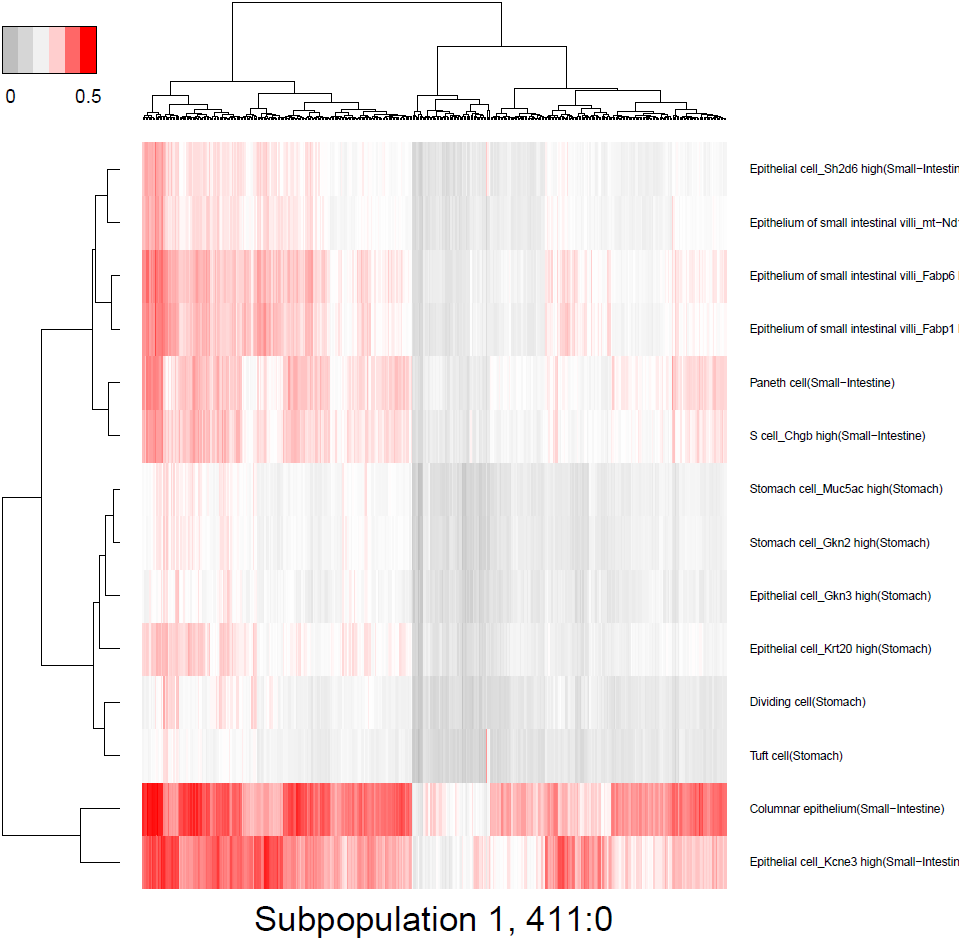
Predicted cell types of the cell subpopulation 1

### 3.3 The cell subpopulation 3

**Figure 7:**
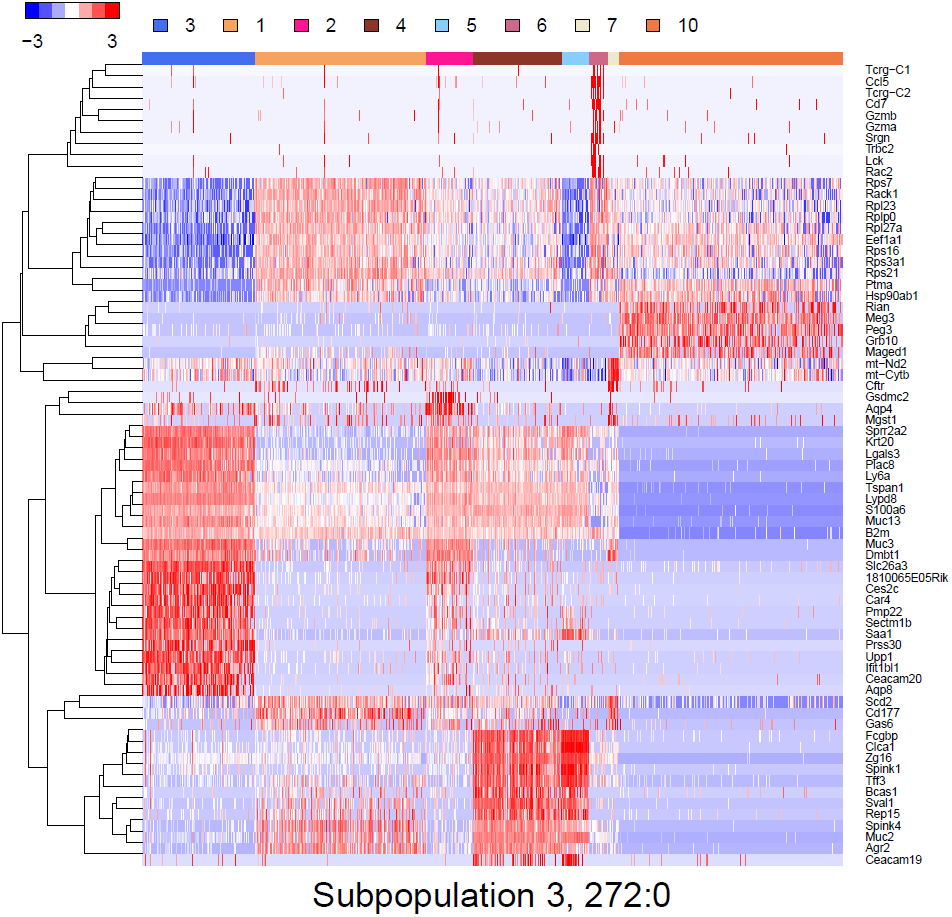
Heatmap of gene expressions that mark the cell subpopulation 3

**Figure 8:**
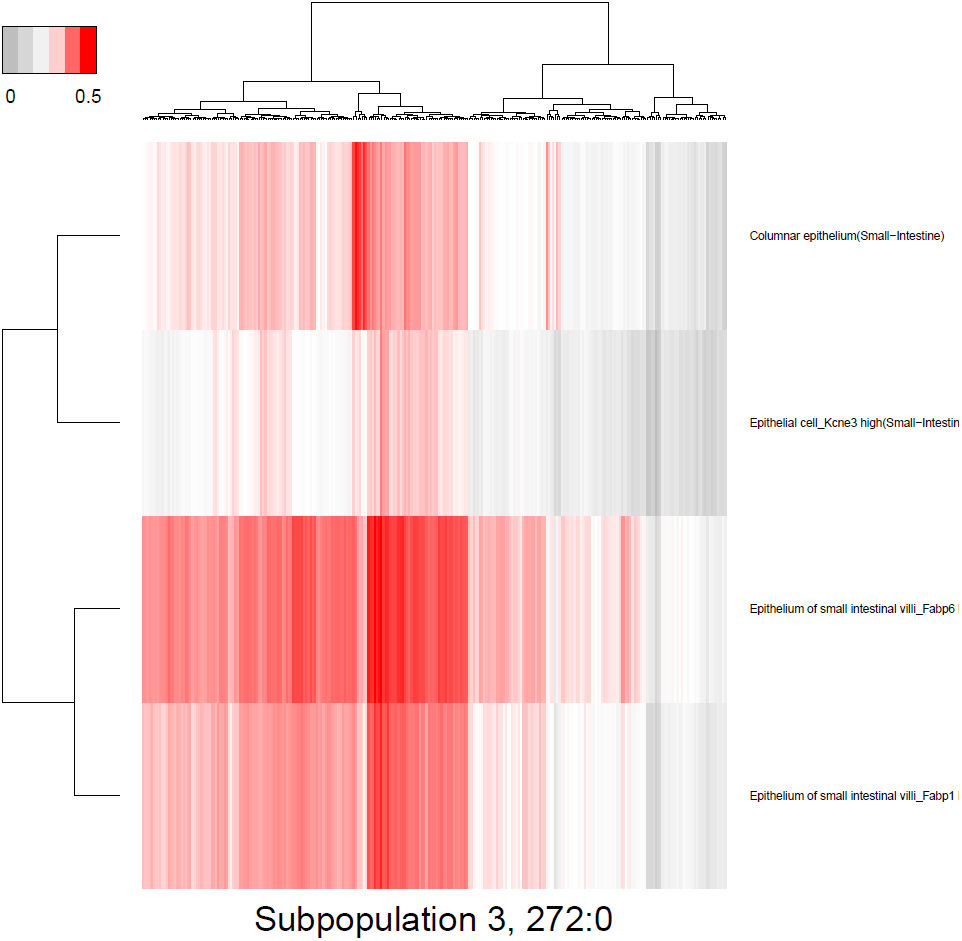
Predicted cell types of the cell subpopulation 3

### 3.4 The cell subpopulation 4

**Figure 9:**
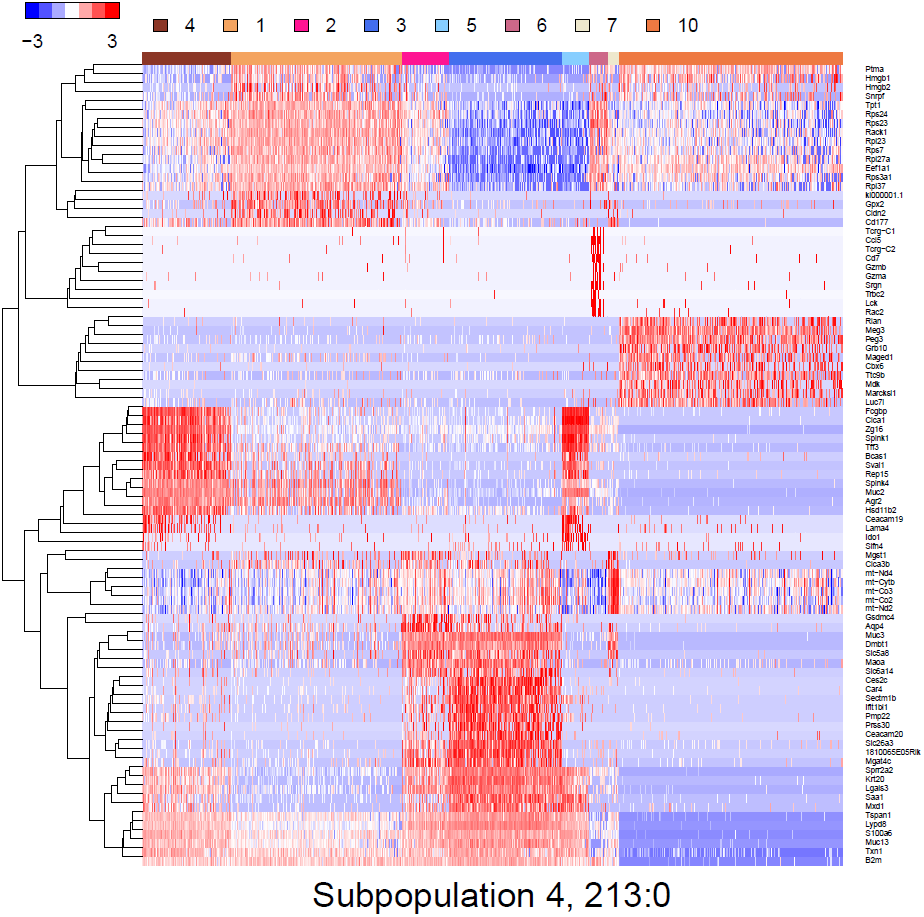
Heatmap of gene expressions that mark the cell subpopulation 4

**Figure 10:**
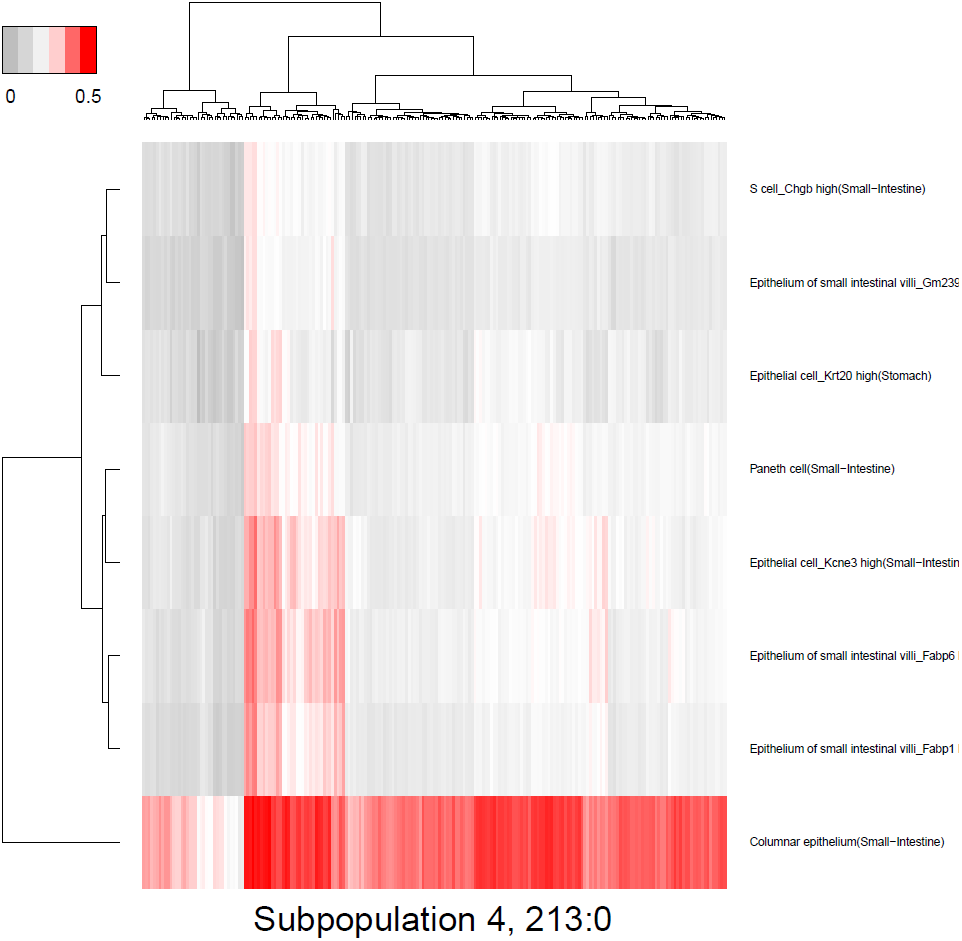
Predicted cell types of the cell subpopulation 4

### 3.5 The cell subpopulation 2

**Figure 11:**
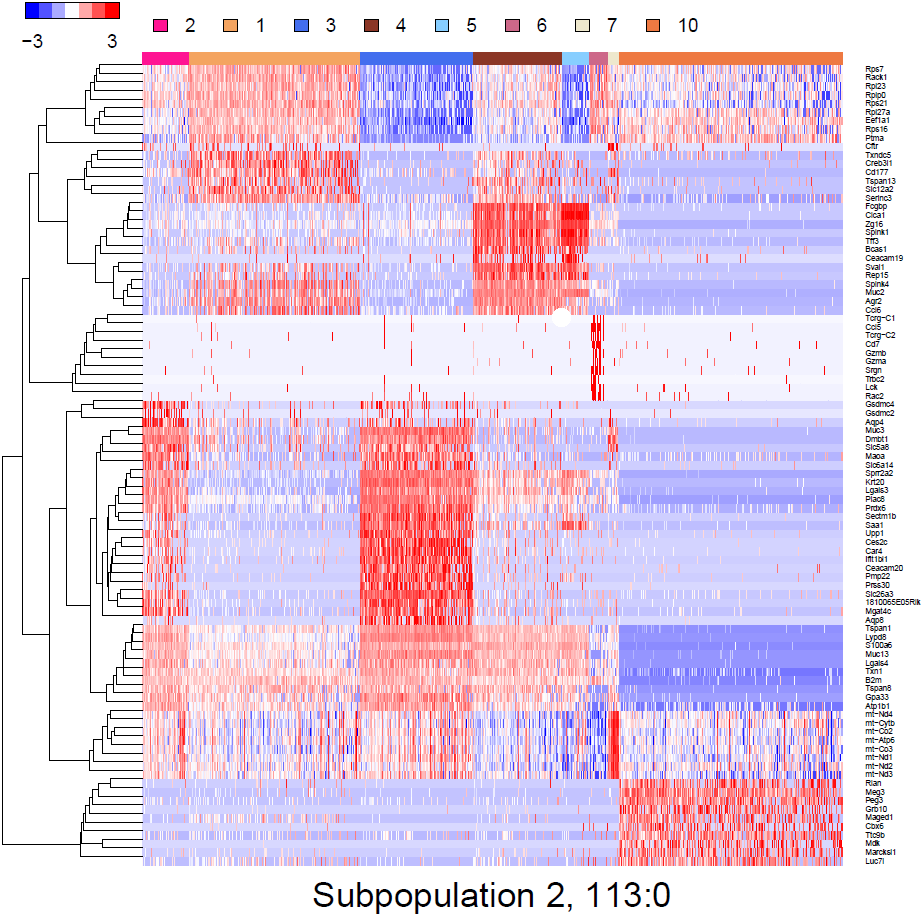
Heatmap of gene expressions that mark the cell subpopulation 2

**Figure 12:**
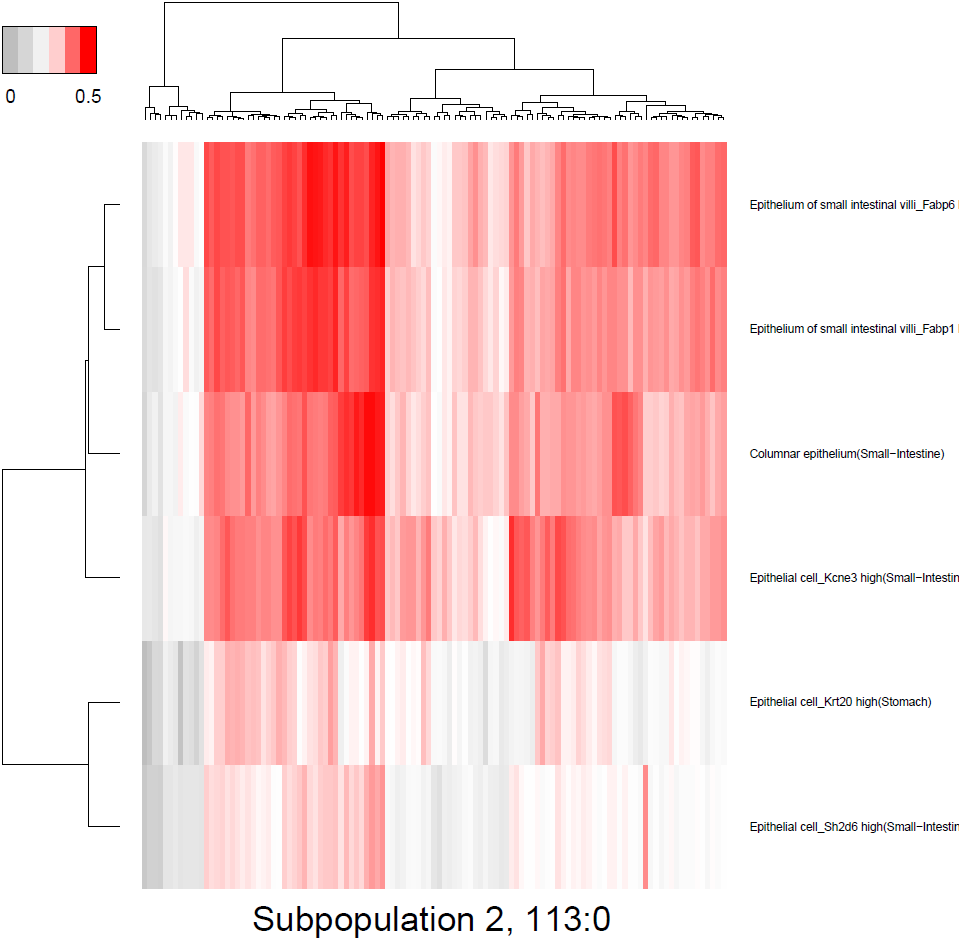
Predicted cell types of the cell subpopulation 2

### 3.6 The cell subpopulation 5

**Figure 13:**
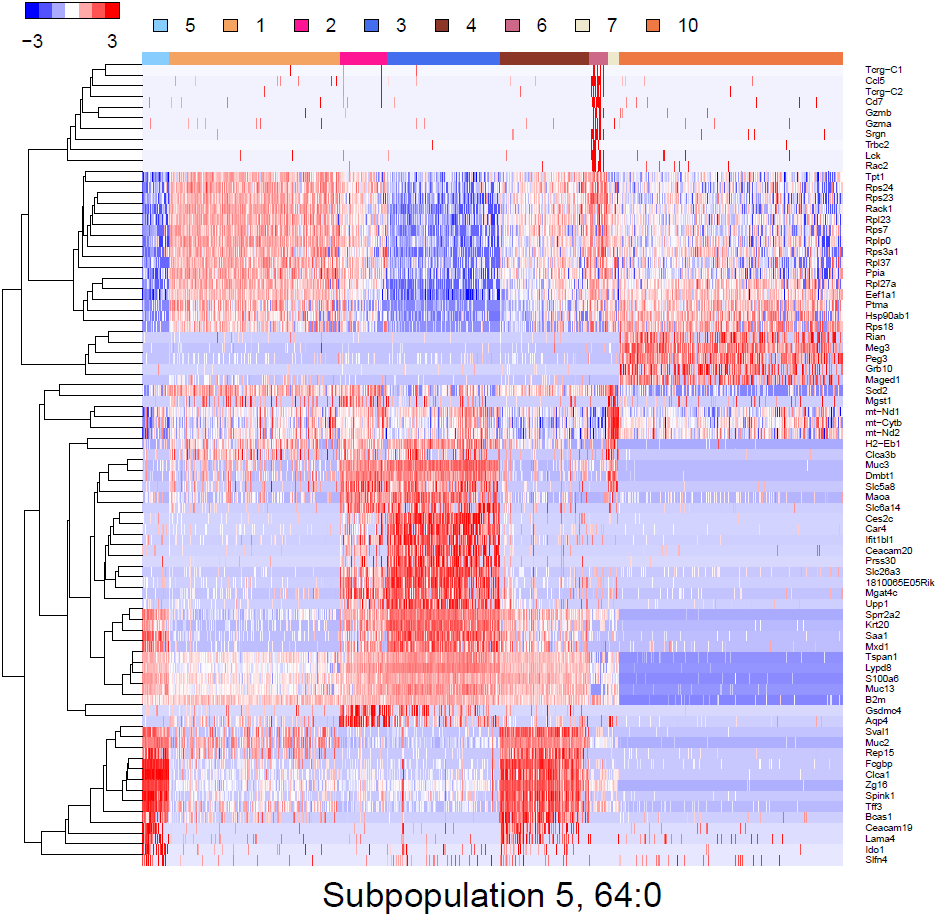
Heatmap of gene expressions that mark the cell subpopulation 5

**Figure 14:**
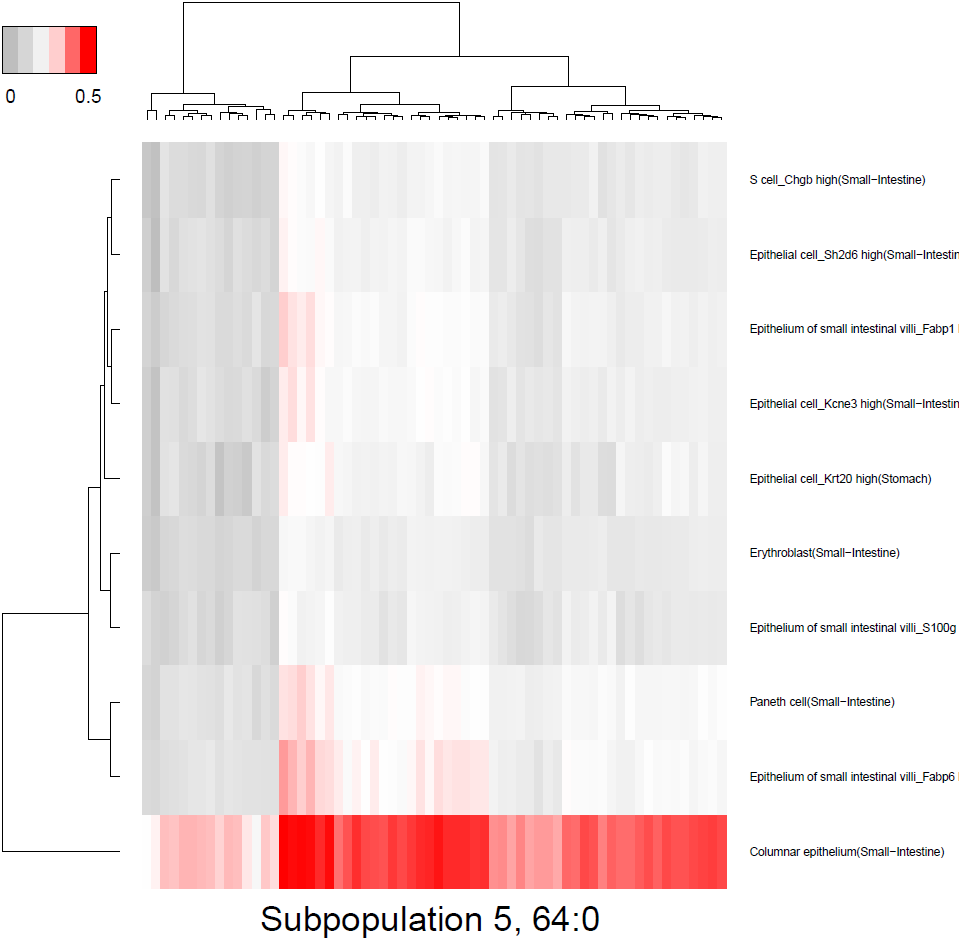
Predicted cell types of the cell subpopulation 5

### 3.7 The cell subpopulation 6

**Figure 15:**
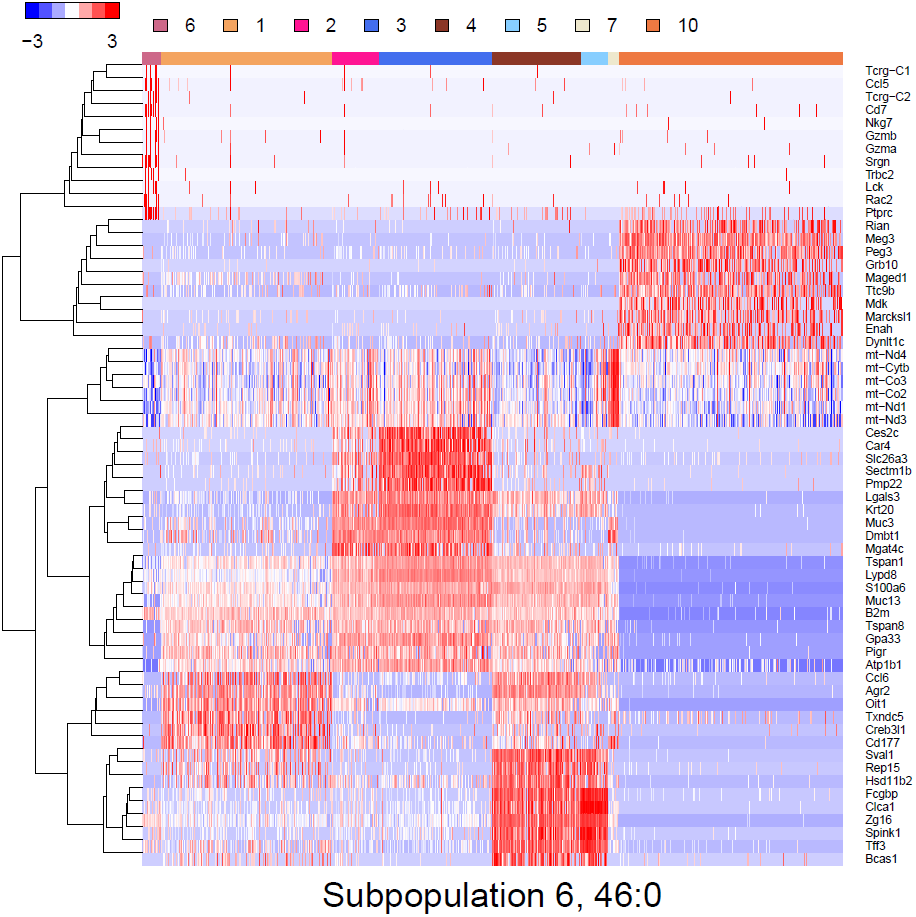
Heatmap of gene expressions that mark the cell subpopulation 6

**Figure 16:**
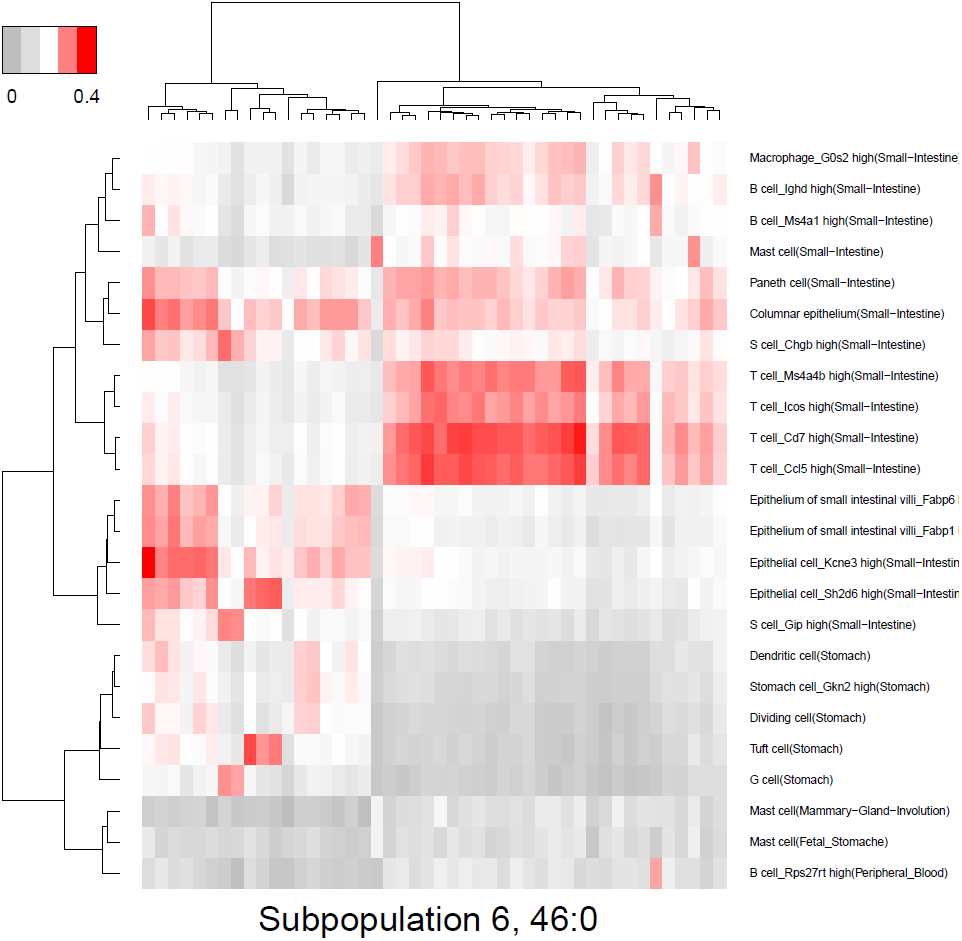
Predicted cell types of the cell subpopulation 6

### 3.8 The cell subpopulation 7

**Figure 17:**
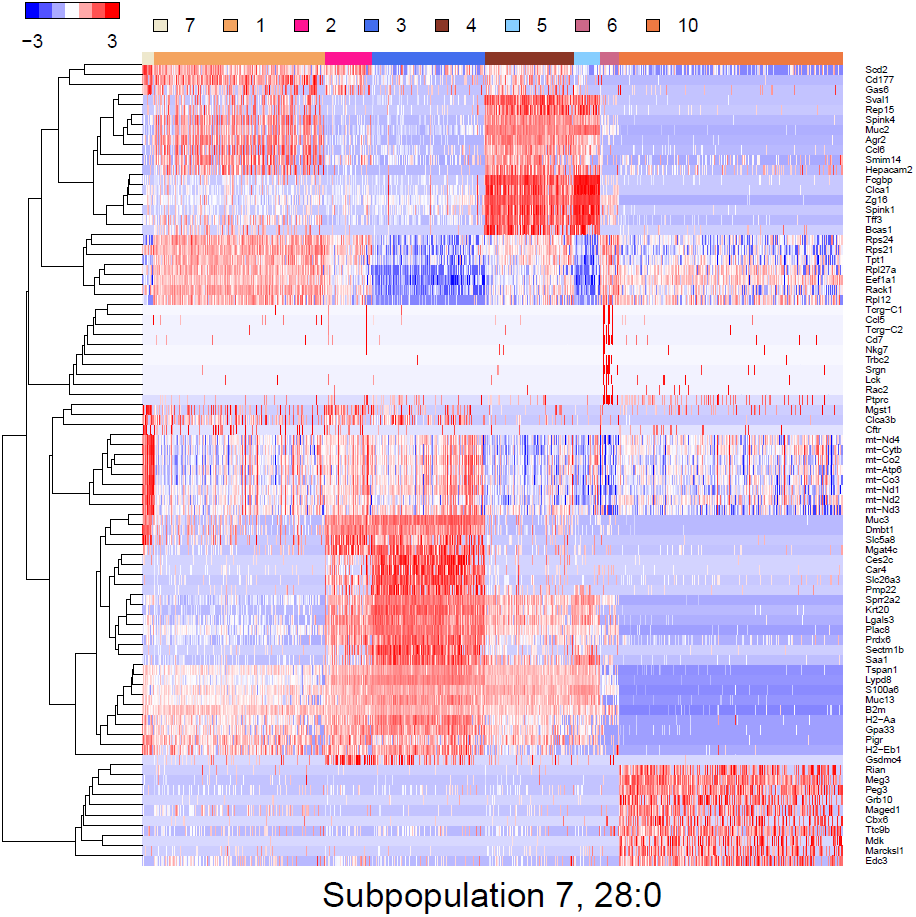
Heatmap of gene expressions that mark the cell subpopulation 7

**Figure 18:**
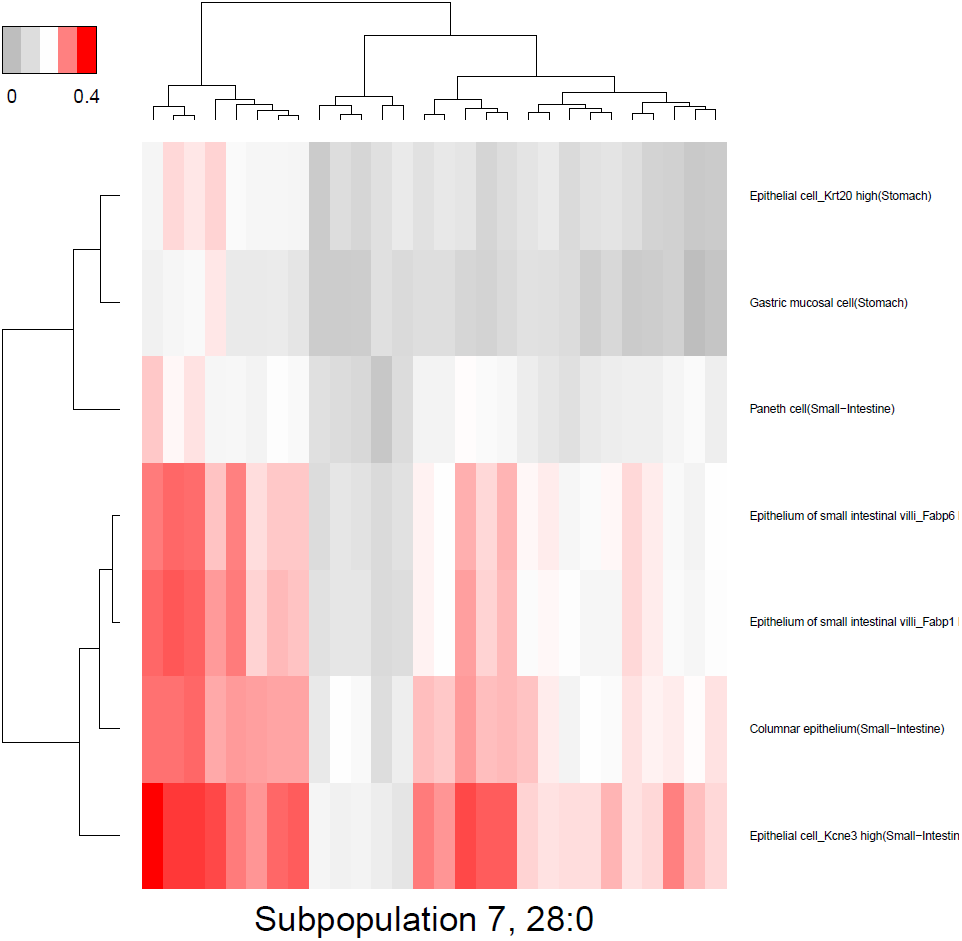
Predicted cell types of the cell subpopulation 7

### 4 Session Info

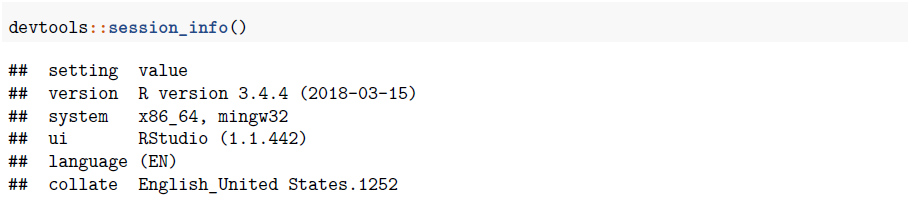
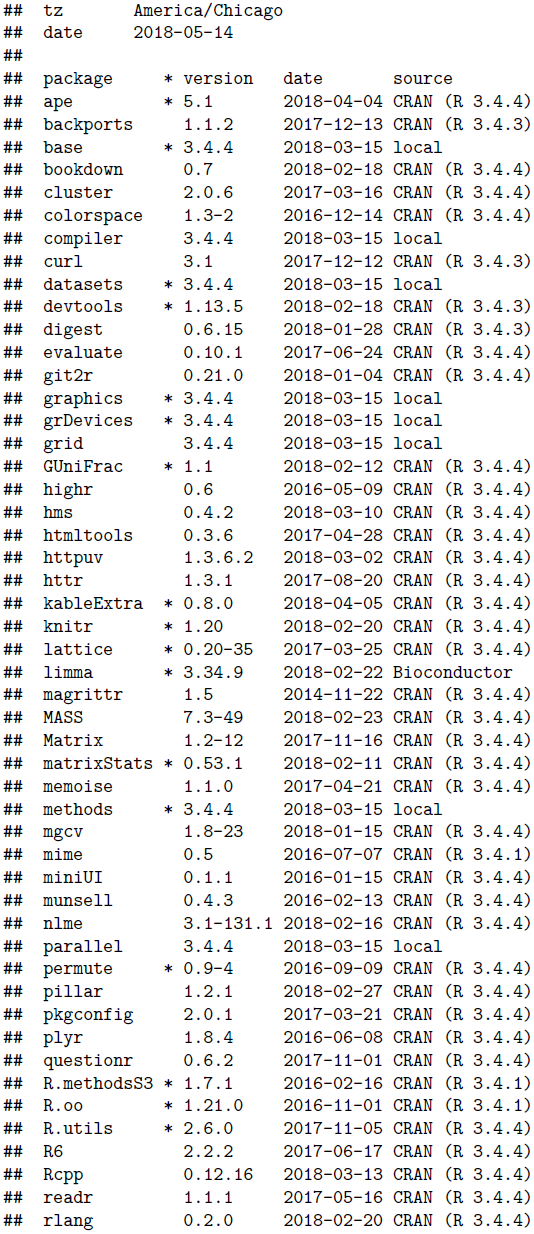
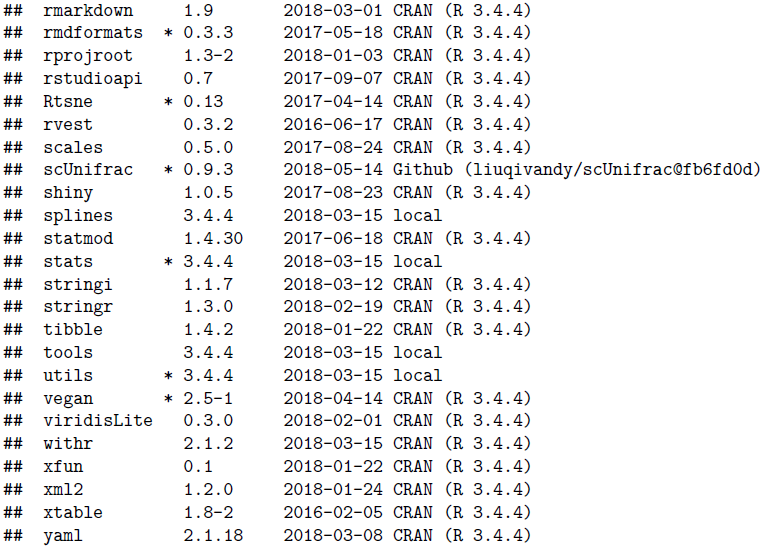

